# Cell type and condition specific functional annotation of schizophrenia associated non-coding genetic variants

**DOI:** 10.1101/2023.06.27.545266

**Authors:** Christine K. Rummel, Miriam Gagliardi, Alexander Herholt, Ruhel Ahmad, Vanessa Murek, Liesa Weigert, Anna Hausruckinger, Susanne Maidl, Laura Jimenez-Barron, Lucia Trastulla, Mathias Eder, Moritz Rossner, Michael J. Ziller

**Affiliations:** Max Planck Institute of Psychiatry, Munich, Germany; International Max Planck Research School for Translational Psychiatry (IMPRS-TP), Munich, Germany; Department of Psychiatry, University of Münster, Münster, Germany; Department of Psychiatry and Psychotherapy, University Hospital, LMU Munich, Munich, Germany; Center for Soft Nanoscience, University of Münster, Münster, Germany

## Abstract

Schizophrenia (SCZ) is a highly polygenic disease and genome wide association studies have identified thousands of genetic variants that are statistically associated with this psychiatric disorder. However, our ability to translate these associations into insights on the disease mechanisms has been challenging since the causal genetic variants, their molecular function and their target genes remain largely unknown. In order to address these questions, we established a functional genomics pipeline in combination with induced pluripotent stem cell technology to functionally characterize ~35,000 non-coding genetic variants associated with schizophrenia along with their target genes. This analysis identified a set of 620 (1.7%) single nucleotide polymorphisms as functional on a molecular level in a highly cell type and condition specific fashion. These results provide a high-resolution map of functional variant-gene combinations and offer comprehensive biological insights into the developmental context and stimulation dependent molecular processes modulated by SCZ associated genetic variation.

## Introduction

One of the most pressing challenges of modern biomedical research is the understanding and treatment of major psychiatric disorders such as schizophrenia (SCZ), bipolar (BD) and major depressive disorder (MDD). Despite decades of intense research, little is known about the molecular mechanisms that contribute to disease onset and progression. In particular, SCZ is characterized by a strong genetic component with a heritability between 75-80%^1,2^, rendering genetics a powerful tool to understand the molecular basis of this devastating disease. Large-scale genome wide association studies (GWAS) have thus far identified 145 regions of the genome that are associated with SCZ at genome wide significance^3^. These loci each harbor hundreds of distinct genetic variants that are all statistically associated with the disease. Furthermore, there are thousands of additional genetic variants that show strong association with SCZ just below the genome-wide significance cutoff, with many likely reaching significance as cohort sizes increase.

However, translating these associations into insights on the disease mechanisms has been challenging due to the fact 90% of the associated genetic variants reside in non-coding regions of the genome with unknown function^4^. Moreover, most of these variants likely act only in disease relevant cell types of the central nervous system, inaccessible to functional experiments. Lastly, each SCZ associated locus can harbor thousands individual disease associated single nucleotide polymorphisms (SNPs) due to linkage disequilibrium (LD), rendering it even more difficult to pinpoint those genetic variants that functionally contribute to the emergence of the phenotype.

Comprehensive functional genomic and epigenomic studies revealed that disease associated common genetic variants are enriched in gene regulatory elements (GREs), in particular in enhancer regions^5,6^. These studies also highlighted that this enrichment is especially pronounced for GREs that are active in cell and tissue types relevant for the respective disease, such as the prefrontal cortex and excitatory neurons in the case of SCZ^7,8^. These findings indicate that a large fraction of non-coding disease associated genetic variants (NCDVs) likely act by modulating tissue/cell type specific gene expression levels, effectively acting as expression quantitative trait loci (eQTLs), contacting their target gene in 3D space as shown by chromatin conformation capture^9,6,17^. This has been further supported by large eQTL studies that map genetic variants and gene expression levels across different cell types and tissues of hundreds of healthy/patient donors^10–12^. Based on these studies, current estimates suggest that between 40% and 50% of SCZ associated SNPs act by modulating gene expression directly in cis^13,14^ with a substantial fraction also altering the chromatin state of associated GREs^12,15^.

While clearly very powerful, contemporary approaches building on epigenomic and qTL information do suffer from multiple critical blind spots: i.) As all these strategies rely purely on profiling and statistical association of varying stringency, they inherently remain correlative in nature. In particular, it remains unclear which of the thousands of NCDVs overlapping with these annotations are indeed functional and which merely associated. Moreover, ii.) currently available epigenomic and qTL datasets at present do not capture some of the most critical SCZ relevant conditions such as many cellular states during human neural development and stimulation dependent conditions. For example, multiple studies in the immune system revealed that 3-18 % of the overall detected eQTL set did not show any effect under baseline conditions, but only contributed to changes in gene regulation upon stimulation with IFNy and *Salmonella*^16^, rendering them stimulation dependent eQTLs. Such observations are particularly relevant in the context of SCZ, which is hypothesized to be at least partially rooted in compromised neurodevelopment and primarily affects neuronal cells, highly responsive to environmental and electrical signals. Thus, for most SCZ associated NCDVs, their cellular and condition specific context of action remains unknown, rendering their biological interpretation at present highly demanding. Lastly, iii.) current annotation information frequently retains a high level of ambiguity, associating individual genetic variants with many putative target genes with entirely distinct biological function within individual GWAS loci (and vice versa). On top of that, many of these genes are expressed in entirely different cellular contexts, rendering the biological interpretation of GWAS hits difficult. Thus, it remains a key challenge to functionally decode the context specific biology of even individual GWAS findings.

In combination, these gaps in our understanding of NCDV biology severely hamper the translation of genetic associations into insights on SCZ related pathomechanisms. This problem is already illustrated by the very basic question on which genetic variant or gene to select for deep functional analysis from the thousands of possible candidates as well as choice of a suitable cellular context.

Here, we set out to overcome these current limitations in the context of SCZ genetics by addressing four key questions: (Q1) Which SCZ associated genetic variants have the capacity to functionally modulate gene expression? (Q2) To what extend is this capacity dependent on cellular context and state? (Q3) What are the target genes of these functional variants in disease relevant cell types? And finally, (Q4) can this annotation be utilized to refine current the biological interpretations of SCZ GWAS?

In order to start to answer these questions, we integrated multiple functional genomics assays (MPRA, CRISPRi, HiC, and single cell technology) into a coherent massively parallel variant annotation pipeline (MVAP, **Figure 1A**), generally applicable to any disease entity with available GWAS data. We then combined this pipeline with disease relevant cellular model systems of the CNS, based on induced pluripotent stem cell (iPSC) and primary mouse cultures to pinpoint the set of molecularly functional SCZ associated genetic variants. These experiments give rise to a unique map of cell and condition specifc NCDVs in SCZ along with their target genes. This map constitutes a valuable resource to guide and enable future experiments to functionally dissect the molecular and cellular mechanisms in SCZ that are driven by these functional NCDVs ^17^. Going beyond this unique resource, our analysis provides several new insights into the condition specific action of NCDVs and their converging effects on key biological pathways in distinct cellular contexts.

**Figure 1.**
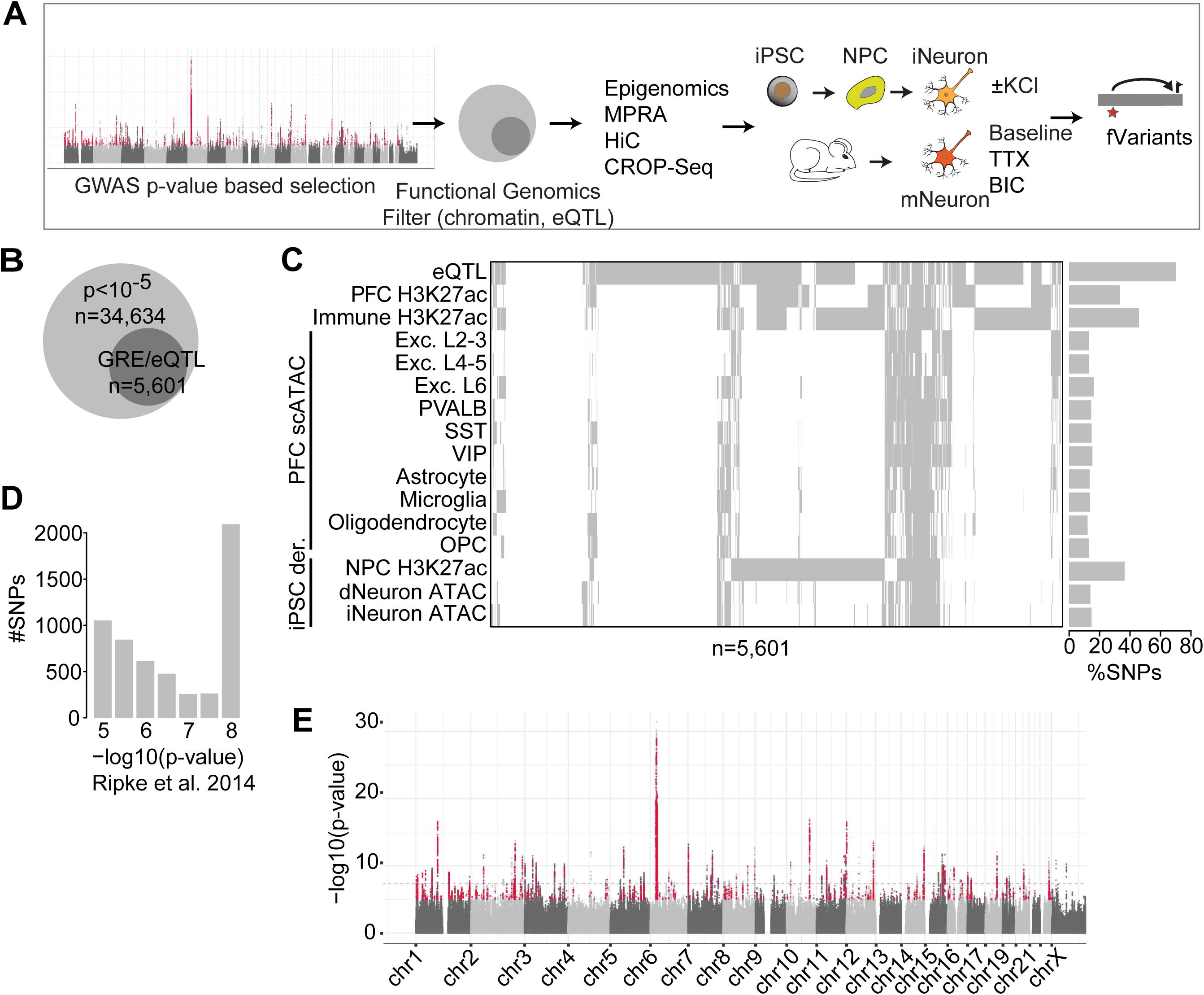
Overview of MVAP pipeline and MPRA design. (A) Schematic of MVAP workflow. (B) All SCZ associated genetic variants below a p-value of 10^−5^ (light grey) and a subset of selected variants for interrogation by MPRA (dark grey) based on epigenomic and eQTL annotation (C) Annotation of all SCZ associated genetic variants selected for MPRA testing (x-axis) with various epigenomic annotations (y-axis) derived from iPSC derived neurons (iNeuron, dNeuron, neural precursor cells (NPCs), single cell ATAC-Seq of adult human post-mortem tissue from the prefrontal cortex (PFC), H3K27ac ChIP-Seq profiles across immune cell types and bulk PFC tissue as well as eQTLs detected in PFC. (D) SCZ association p-value distribution of selected SNPs based on Ripke et al. 2014. (E). Manhattan plot showing all (grey) and for further investigation selected (red) schizophrenia associated SNPs across the genome. Blue dotted line depicts GWAS significance cutoff at p-value<10^−8^.

## Results

### Identification of candidate expression modulating genetic variants associated with SCZ

In order to identify SCZ associated functional genetic variants (Q1), we first sought to pinpoint those SNPs that have the capacity to modulate gene expression levels in an allele specific manner, termed expression modulating variants (emVars)^18^. Given that genetic variants with strong, but sub-threshold association signal that overlap with GREs are frequently molecularly functional and often achieve genome wide significance with increasing GWAS cohort sizes, we decided to also interrogate the latter variants set ^19^.

To that end, we annotated all SCZ associated SNPs below a p-value of 10^−5^ based on the PGC GWAS from 2014 ^20^ with a panel of publicly available and newly generated epigenomic and eQTL data from disease relevant cell types (**Table S1**) in order to filter the set of candidate variants. This analysis identified 5,601 SNPs overlapping (**Figure 1B**) with at least one annotation and fulfilling our selection criteria (**Figure 1C**, see **Methods** for details). This selected set of candidate variants included a large fraction of SNPs that achieved genome wide significance, but is also composed of more than 50% sub-threshold variants (p-value higher than 5×10^−8^) (**Figure 1D**). Together, these variants were distributed across the entire genome, covering all genome wide-significant loci (**Figure 1E**) and overlapping with annotation for eQTLs in human adult prefrontal cortex (PFC) tissue (80%), chromatin marks for active enhancers in the post mortem PFC (~50%) as well as multiple other cell type specific open chromatin regions (**Figure 1C**).

### Assessing the reproducibility of massively parallel reporter assays in disease relevant cell types

In order to interrogate this variant set for their cell type specific capacity to modulate gene expression levels, we generated massively parallel reporter assay (MPRA) libraries harboring SCZ associated SNPs. This assay allowed us to functionally test thousands of individual putative regulatory elements for enhancer activity (referred to as putative enhancer element, pEEs) as well as the impact of single nucleotide variants on pEE function. Briefly, thousands candidate pEEs are synthesized using DNA synthesis technology, each in two versions: harboring the SCZ associated reference or alternative allele (referred to as allele specific pEEs, **Figure S1A**). Subsequently, all pEEs are cloned as a pool 5’ to a minimal promoter followed by a reporter and a unique 16mer barcode into a lentiviral plasmid library^21^. Each allele specific pEE is associated with multiple individual barcodes (**Figure S1E**). Once transduced into the cells of interest, next-generation sequencing of the mRNA fraction originating from the integrated lentiviral reporter constructs then give rise to a digital measure of the individual reporter construct activity by counting the number of observed 3’ barcode tags associated with each pEE reporter construct. The comparison of the pEEs’ activity with either allele allows us to determine whether or not the SCZ associated SNP has the potential to modulate gene expression levels within the context of the reporter assay.

Here, we independently synthesized three MPRA library pools, given the large number of individual genetic variants to be tested and constraints on the available cell numbers of CNS cell types. More specifically, we generated one smaller, developmentally focused MPRA library, containing many variants overlapping with putative enhancer regions in iPSC derived NPCs. This developmental library (DevLib) contained in total 5,118 distinct reporter constructs, assaying 2,559 distinct SNPs with two alleles each, as well 85 positive control constructs and 152 negative control constructs. In addition, we also generated a larger MPRA library focused on SNPs overlapping with post mortem eQTLs and open chromatin regions in adult post mortem prefrontal cortex (eQTLLib), assaying 4,325 distinct SNPs (8,650 allele specific pEEs). Lastly, we generated a focused validation library of high complexity that included 215 SCZ associated SNPs from the DevLib as well as 100 random SNPs not associated with SCZ as additional controls (**Table S2**).

Each library contained between 370,705 and 927,647 individual barcodes (**Figure S1B-D**), translating into a median individual barcode coverage of 29, 45, and 500 per tested allele specific pEE (**Figure S1E**). Between 60-95% of the originally selected SNPs for allele specific analysis were recovered (see **Methods**, **Figure S1F**). This allowed us to assess in total 9,902 allele specific pEEs representing 4,951 pEEs.

Subsequently, we evaluated the resulting MPRA libraries across a panel of disease relevant conditions via lentiviral transduction in iPSC derived neural precursor cells (NPCs), iPSC derived neurons (iNeurons) under baseline and stimulation conditions (Baseline/Stimulation) as well as primary cultures of mouse cortical neurons at DIV 12 under baseline, stimulation (Stimulation) and tetrodotoxin (TTX) conditions with 3-5 replicates per condition (**Table S3, Figure S2**). Prior to further analysis, the allele specific pEE activity was normalized to its abundance in the original plasmid library (**Figure S1G**). Overall, the MPRA experiments show high reproducibility across conditions (**Figure 2A**, **Figures S3A,B**), with reproducibility depending on individual library complexity and the abundance of each element within a library (**Figure 2A**, **Figures S3A-C**).

**Figure 2.**
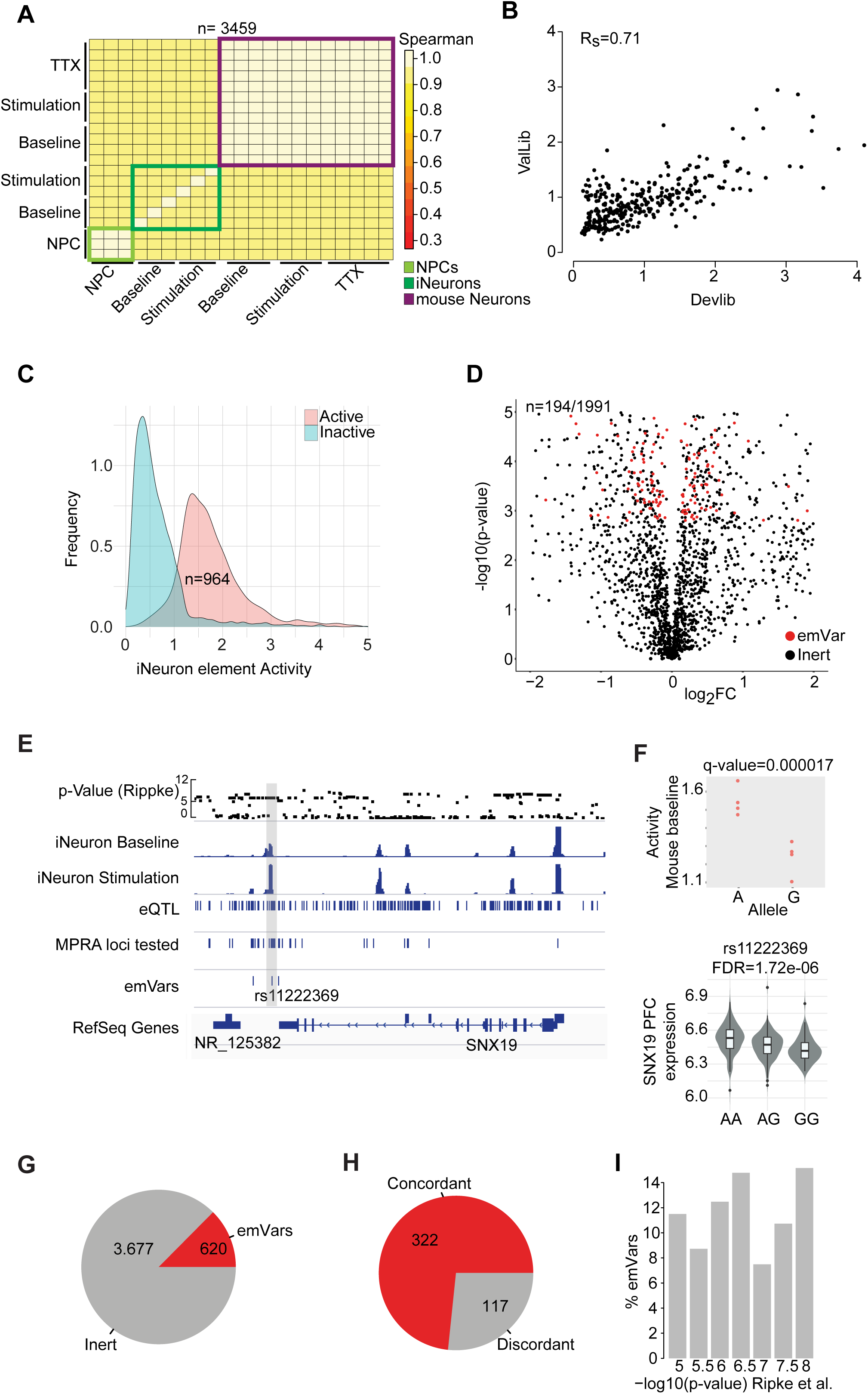
Reproducibility and differential allele activity of SCZ associated genetic variants. (A) Spearman correlation heatmap of cpm normalized log2 ratio between the MPRA mRNA counts and their abundance in the original plasmid library for all elements measured in all conditions of the DevLib. (B) Consistency of sequence element activity contained in both the DevLib and the ValLib (n=193) in mouse primary neuronal cultures under baseline conditions. (C) Distribution of normalized pEE activity (alpha-value, x-axis) in iNeurons for elements delineated as active (red) or inactive (blue) based on comparison to negative control elements (p-value ≤0.005). (D) Volcano plot of p-value (-log10, y-axis) and log_2_ fold change between all SCZ associated allele specific pEE evaluated in iNeurons in the DevLib (n=1,991). Elements with allele specific activity are shown in red (emVars, FDR≤0.005 and minimal activity p-value≤0.005). (E) Example IGV plot of the SNX19 locus depicting from top to bottom: the GWAS based p-value of SCZ association for all genetic variants, the ATAC-Seq profiles of iNeurons under baseline and stimulated condition, the location of eQTLs in adult human post mortem PFC tissue, the location of all SNPs interrogated by MPRA, and all SNPs associated with allele specific GRE activity (emVars). (F) Top: Allele specific GRE activity for the A and G allele of the rs11222369 SNP also shown in c. based on MPRA measurements in primary mouse neuronal cultures under baseline conditions. Each dot represents an independent MPRA assay replicate. Bottom: Expression of the SNX19 gene in human PFC by donor allele status of the rs11222369 eQTL (FDR=1.72e-06). (G) Fraction of all SCZ associated genetic variants successfully interrogated by MPRA (grey) and identified as emVars (red) across all MPRA experiments. (H) Fraction of emVars overlapping with eQTLs in PFC that show significant concordance or discordance with respect to the allele specific up/down-regulation of the overlapping eQTL in at least one tested cell type/condition red or grey, respectively. (I) Percentage of all emVars (y-axis) as a function of SCZ association p-value based on the Ripke et al. 2014 GWAS.

We reasoned that the most robust method to validate observations from the larger MPRA libraries would be through the generation of a smaller, but high complexity validation library, comprising a subset of originally tested variants (ValLib, **Figure S3D**). We thus first leveraged this high complexity library to assess the correlation with each larger discovery MPRA library and found good agreement between the allele specific measurements on the joint SNP sets (Spearman correlation 0.71, **Figure 2B**) as well as between the DevLib and eQTLLib (0.73 **Figure S3E**). As expected, average Spearman correlation increased with higher pEE abundance (**Figure S3F**).

### Identification of SCZ associated emVars using MPRA

In the next step, we determined the set of pEEs with the capacity to operate as an enhancer element (EE), leveraging the negative control elements as reference (FDR ≤ 0.005, **Figure 2C**). Subsequently, we only considered these active regions in the context of allele specific analyses to identify those EEs where a change in allele specific activity would be meaningful. In order to further calibrate the thresholding parameters to define regions with allele specific activity, we carried out extensive sensitivity and specificity analyses using the high complexity validation library as well as our set of negative control SNPs as reference (**Figures S3G,H, Methods**).

Based on these analyses, we selected thresholding parameters (FDR ≤ 0.005 and activity threshold p-value ≤ 0.005) that allowed us to assess allele specific EE activity with a TPR of 55% and a FPR of 9%, providing a compromise between sensitivity and specificity. Importantly, this approach resulted in active enhancers that were enriched for location in open chromatin marks and eQTLs in relevant cell types **(Figure S3I)**. Application of this analysis strategy to the iNeuron condition (DevLib), identified 194 emVars (of 1991 tested pEE) (**Figure 2D**). We note, that there is a substantial fraction of SNPs (675) that show evidence for allele specific activity, but are located in elements below the minimal activity level threshold (referred to as non-enhancer emVars). The set of emVars included several variants at the well-known SCZ risk locus *SNX19* (**Figure 2E**), with the highest confidence variant rs11222369 showing robustly reduced MPRA activity for the G allele in the mouse baseline condition (**Figure 2F top**). This reduction in EE activity is in line with publicly available human post mortem eQTL data for the PFC, revealing significantly reduced expression of the *SNX19* gene for the GG genotype (**Figure 2F bottom**).

In total, we identified 620 emVars (12.5% of the variants tested by MPRA, 1.7% of the originally considered SNPs) across the union of the DevLib and eQTLLib and all tested conditions (**Figure 2G**), similar to the fraction of functional variants observed in previous MPRA experiments in other systems^18^. Validation of 44 emVars using the separate high complexity ValLib confirmed 65.9% fulfilling both minimal activity and allele specific activity thresholds in the ValLib as well as 86% fulfilling only the allele specific activity threshold (non-enhancer emVars). Moreover, almost 75% of the identified emVars that overlapped with an eQTL in PFC data showed concordance in the direction of the allelic effect (**Figure 2H**). These observations are consistent with previous findings of MPRA experiments in K562 cell^18^ and provide further support for the relevance of the emVars defined by MPRA. Interestingly, the identified emVars were distributed equally across the entire p-value spectrum of SCZ association (**Figure 2I)**

### Cell type and condition specificity of emVars

Based on the notion that GREs as well as eQTLs operate in a highly cell type specific fashion, we next sought to determine to what extend EE as well as emVar activity was cell and condition specific (Q2). To that end, we evaluated the allele specific activity of SCZ associated EEs across a range of distinct cellular states and conditions using MPRA. In particular, we investigated the developmental specificity of emVar activity in iPSC derived NPCs and iPSC derived mature neurons. This setup enabled us to specifically test the hypothesis whether or not a subset of SCZ associated variants operate at early developmental time points and thus are more likely to contribute to the developmental component of SCZ. This approach identified almost half of all tested allele specific EEs to operate in a cell type specific fashion (**Figure 3A**), consistent with their chromatin state pattern by which they were selected (**Figure 3B**). Moreover, a large fraction of EEs showed differential activity between the two developmental conditions for one or both alleles (**Figure 3C**). This specificity in activity pattern directly translated into the cell type specific differential allele activity, with 40% of emVars showing allele specific activity only in NPCs or iNeurons (**Figure 3D**, **Figures S4A, B**). The set of cell type specific emVars included a SCZ associated variant (rs9806806) at the NMDA receptor subunit gene *GRIN2A*, not previously identified as an eQTL (**Figure 3E**). This variant exhibited highly significant differential activity in iNeurons (FDR 0.0002), while the observed difference in NPCs was far less pronounced (**Figure 3F**), consistent with expression of *GRIN2A* only in more mature neuronal cells.

**Figure 3.**
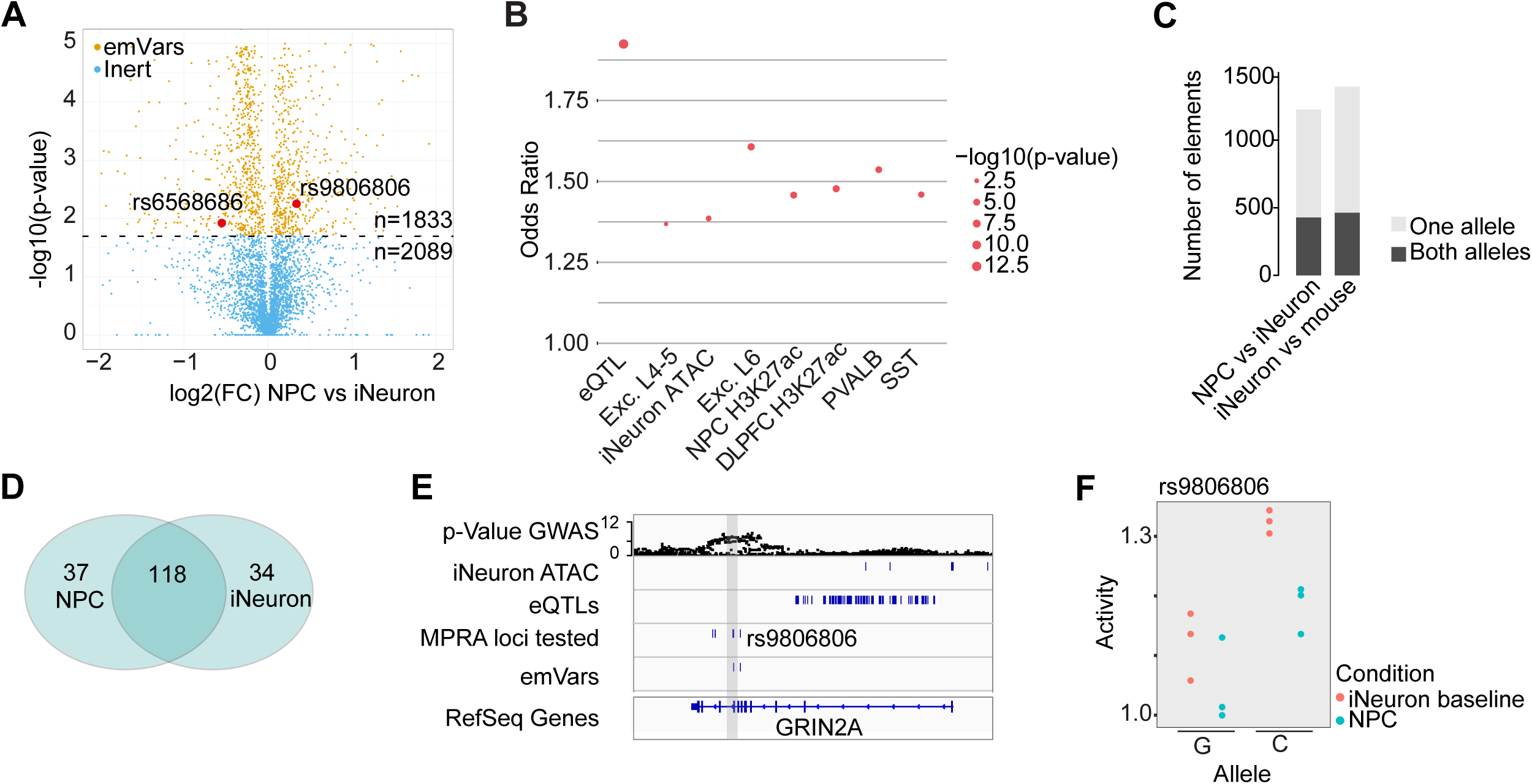
Developmental stage specific GRE and emVar activity. (A) Comparison of allele specific EE activity of the DevLib MPRA library between NPCs and iPSC derived iNeurons at day 49. Y-axis depicts-log10 p-value and x-axis log2 fold change of the comparison: negative values indicate higher activity in the NPCs. Yellow dots indicate emVars with significantly differential activity between the conditions (FDR≤0.05). (B) Enrichment of emVars differentially active between NPCs and iNeurons overlapping with distinct genomic features, including: eQTL – eQTL (FDR≤0.05) in adult human post mortem PFC, single cell ATAC-Seq in PFC: Exc. L4-5 – excitatory neurons layer 4-5, Exc. L6 – excitatory neurons layer 6, PVALB – parvalbuminergic interneurons, SST – somatostatin interneurons, NPC H3K27ac – ChIP-Seq for H3K27ac in iPSC derived NPCs, PFC H3K27ac – ChIP-Seq in dorso lateral prefrontal cortex of adult human post mortem brain, iNeuron – ATAC-Seq iPSC derived neurons at day 49 of culture. y-axis indicates odds ratio and point size p-value of Fisher’s-exact test results. (C) Number of DevLib MPRA EE (y-axis) where both interrogated alleles (dark grey) or only one allele (light grey) showed differential activity between the conditions indicated on the x-axis in mouse baseline. (D) Overlap of detected emVars in NPCs and iNeurons. (E) IGV annotation of an example SCZ associated emVar detected at the GRIN2A locus. (F) Allele specific GRE activity for the G and C allele of the rs9806806 SNP based on MPRA measurements in NPCs (blue) or iNeurons (red). Each dot represents an independent MPRA assay replicate.

This cell type and condition specificity of emVars was not limited to NPCs and iNeurons, but was also observed between iNeurons and mouse neurons (comprising different neuronal cell types, e.g. also inhibitory neurons, **Figure S4A, B**) and distinct neuronal activity states (e.g. baseline, stimulated, and TTX) (**Figures 4A,B**). Consistent with their selection, most emVars were enriched in the open chromatin regions corresponding to the condition in which they were identified (**Figure 4C**). However, several emVar sets were also enriched in open chromatin regions from other cell types (**Figure 4C**). This observation is consistent with a substantial number of emVars showing allele specific activity in a pan-neuronal manner (**Figure 4B**). Importantly, the latter emVar set was located in EEs that were H3K27ac positive across several other cell types and tissues (including many non-neuronal) compared to emVars with more restricted cell/condition specificity (**Figure S4C**). Moreover, these observations also highlight the utility of primary mouse neuronal cultures, suggesting that the overall trans-acting machinery driving EE and emVar activity is conserved between the species, despite the fact that the SNPs and EEs themselves are not present in the mouse genome.

**Figure 4.**
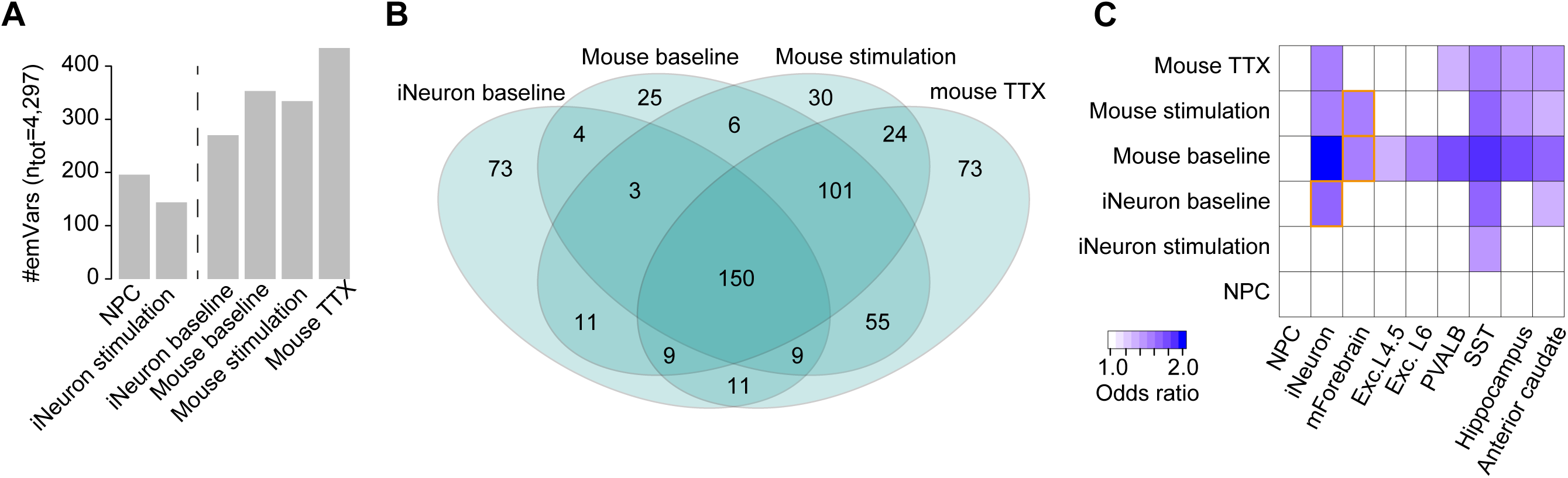
Summary of emVars across conditions. (A) Number of emVars detected across all measured conditions combining Dev and eQTL MPRA library. Dashed line separates conditions only measured for the Dev Lib (left) and conditions measure for all investigated MPRA libraries (right). (B) Overlap of emVars detected across cell types and conditions measured for both Dev and eQTL lib. (C) Enrichment of emVars compared to non-emVars overlapping with distinct genomic features (eQTLs show no enrichment, not shown), including: NPC-H3K27ac ChIP-Seq for NPCs, iNeurons - ATAC-Seq in iNeurons, mForebrain ATAC-Seq in primary mouse forebrain at passage 0, single cell ATAC-Seq in PFC: Excl.4-5 – excitatory neurons layer 4-5, Exc. L6 – excitatory neurons layer 6, PVALB – parvalbuminergic interneurons, SST – somatostatin interneurons, Hippocampus/anterior caudate: ChIP-Seq for H3K27ac in human post mortem brain tissue. Color indicates odds ratio based on Fisher’s exact test, only significant enrichments (p-value≤0.05). Orange boxes indicate matching conditions between MPRA experiment and ChIP-Seq/ATAC experiments.

Neurons do not only alter their transcriptional and epigenetic landscape as a function of cellular identity or developmental state, but also in response to external stimuli. In fact, mature neurons are highly plastic cell types that adjust key components of their gene regulatory program in response to a complex code of stimuli, ranging from neurotransmitter induced depolarization to hormone mediated long lasting signals. These input signals critically determine the response of the neuron to future stimuli, in particular in the context of learning processes that involve various types of stimulus dependent GREs. Therefore, we hypothesized that the response strength and regulatory capacity of stimulus dependent EEs are susceptible to genetic alterations. Against this background and the importance of stimulus dependent processes for neuronal plasticity and mental illness, we next determined to what extend SCZ associated genetic variation might exert its effect in a neuronal stimulus dependent manner. To that end, we determined the overall activity of the eQTL and DevLib in electrophysiologically highly active iNeurons (**Figures S5A-C**) under stimulation (high potassium containing media which depolarizes the entire neuron), as well as in primary mouse neurons derived from the cortex under bicuculline/4-AP/glycine/strychnine (inhibiting inhibitory postsynaptic input, therefore indirectly stimulating the neuronal network) or TTX (disrupting action potential generation) treatment. Comparison of EE activity under stimulation and TTX (representing the greatest contrast) revealed widespread stimulus dependent activity (**Figure 5A**).The vast majority of the EE exhibited higher activity under stimulation conditions, consistent with previous reports based on epigenomic profiling^22,23^. Stimulus responsive EE included many open chromatin regions detected in human post mortem brain (**Figure S5D**) and validated a SCZ associated region in the *CACNA1C* gene (**Figure 5B**). In line with its activation upon neuronal stimulation, the latter GRE contained a binding site for the classic activity dependent trans-acting factor CREB (**Figure 5B bottom**). The presence of transcription factor binding sites for stimulus dependent trans-acting factors was widespread among stimulus responsive EEs, with many well-known activity dependent transcription factor binding sites of CREB, ETV, ATF and AP-1 transcription factors ^24^ being enriched with respect to the non-responsive set (**Figure 5C**).

**Figure 5.**
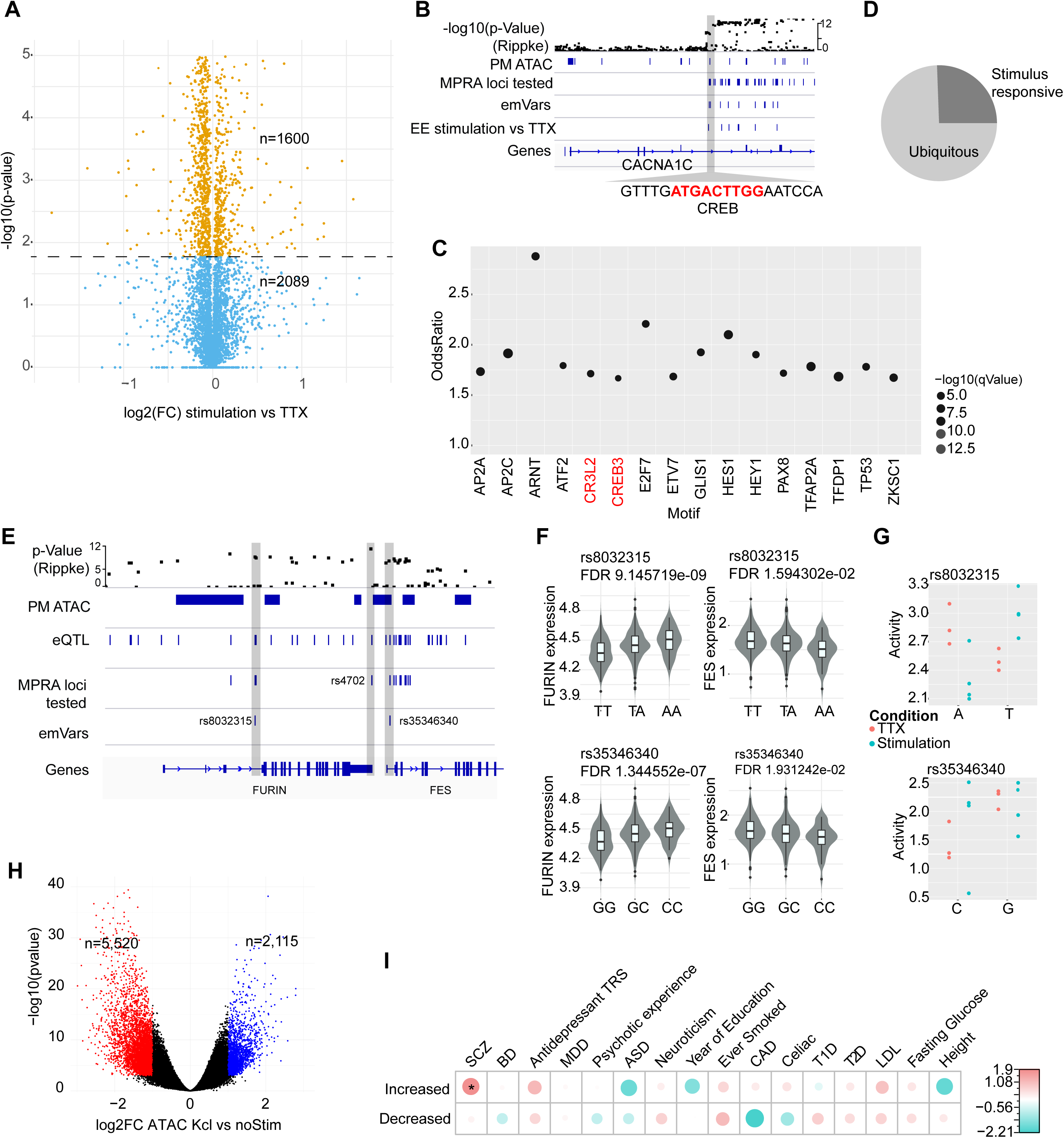
Characterization of neuronal activity dependent SCZ associated eQTLs. (A) Comparison of EE activity of the DevLib MPRA library between primary mouse cortical cultures with abolished (TTX) and stimulated electrophysiological network activity (Stimulation, inhibiting interneuron driven neuronal neurotransmission, leading to hyperactivity). Y-axis shows −log10(p-value) and x-axis log2(fold change) of the comparison, where negative values indicate higher activity in the BIC condition. Yellow dots indicate sequence elements with significantly differential activity between the conditions (FDR≤0.05). (B) Example IGV characterization of stimulation dependent sequence elements at the CACNA1C locus (grey box). Annotation includes GWAS based p-value of SCZ association, pm ATAC-post mortem scATAC peaks, MPRA loci tested – all sequence elements interrogated by MPRA, emVars – all SCZ associated variants with significant allele specific activity; BIC vs TTX – all sequence elements with significant differential activity between mouse primary cortical cultures under stimulation and TTX treatment. Grey triangle shows a high-resolution view of a part of the sequence element highlighting the presence of a CREB binding motif. (C) Distribution of transcription factor motif (x-axis) enrichment (y-axis, odds ratio Fisher’s exact test) in stimulation responsive sequence elements compared to non-stimulation responsive based on mouse MPRA experiments. (D) Fraction of stimulation dependent emVars (stimEmVars, dark grey) relative to all emVars, requiring that allele specific activity was only observed in cell culture conditions (iNeurons or primary mouse neurons) exhibiting electrophysiological activity (stimulation, baseline) but not under TTX treatment. (E) Example of two stimEmVars at the FURIN/FES locus, grey boxes indicate location of stimEmVars. (F) Expression of the FURIN (left) or FES (right) gene in human post mortem PFC by donor allele status (n=468 donors) of the rs8032315 (top) or rs35346340 (bottom) eQTL for each gene. (G) Allele specific GRE activity for the A and T allele of the rs8032315 (top) or the C/G allele of the rs35346340 (bottom) SNP based MPRA measurements in primary mouse neuronal cultures under stimulation (blue) or TTX (red) conditions. Each dot represents an independent MPRA assay replicate. (H) Differential peak accessibility between baseline and stimulation condition iPSC derived iNeurons at day 49 measured by ATAC-Seq across n=10 distinct donors per treatment conditions. Results are shown by p-value (y-axis) and log2 fold change per peak, where red dots indicate peaks with increased accessibility upon stimulation (n=5,520), blue decreased accessibility (n=2,115) and black unchanged (n=228,841). (I) Cell type specific partition heritability analysis for all peaks with increased or decreased accessibility by ATAC-Seq upon stimulation of iPSC derived iNeurons across a panel of traits based on GWAS summary statistics, including BD – bipolar disorder, Antidepressant TRS – treatment resistant depression, MDD – major depression, ASD – autism spectrum disorder, CAD – coronary artery disease, T1D/T2D – Type 1/2 diabetes, LDL – Levels of low-density lipoprotein. Dot size indicates enrichment levels as z-scores and * indicates significance after Benjamini-Hochberg correction (FDR≤0.05).

Based on these results, we next assessed the presence of allele specific response to stimulation, comparing conditions of various stimulation intensity (baseline, depolarization or disinhibition stimulation) against the TTX condition, where inter-neuronal communication was inhibited. This analysis revealed that ~25% of emVars indeed operate in a stimulation dependent manner (**Figure 5D**). Similar to all stimulus responsive elements, transcription factor motifs of canonical stimulus dependent trans-acting factors, including EGR, NR4A1/2 and REST were overlapping precisely with stimulus dependent emVars (stimEmVars).

The set of stimEmVars included two SCZ associated SNPs at the well-known SCZ associated locus FURIN/FES (**Figure 5E**), which has also been associated with various other diseases of the CNS^25^ and other organs^26^. Importantly, both emVars have been annotated as potent eQTLs with opposite effects on both genes (**Figure 5F**). Interestingly, rs8032315 showed higher activity for the T allele under TTX conditions (A>T, **Figure 5G** red top), while this effect switched under stimulation (T>A, **Figure 5G** turqise top). Moreover, the latter allele specific activity pattern was consistent with the eQTL effect of this particular SNP on the FURIN gene (**Figure 5F** top left), while the former is consistent with the eQTL effect on the FES gene (**Figure 5F** top right).

Moreover, one of the identified stimEmVars was in close proximity (1.3kb) to a previously identified eQTL (rs4702) of the *FURIN* gene with profound effects on cell physiology in iPSC derived neurons ^17^. Consistent with the latter study, our MRPA based test of the rs4702 variant revealed highly allele specific activity in iNeurons and mouse primary culture (FDR=0.000259), but it was classified as a non-enhancer emVar. This observation is consistent with the location of the rs4702 variant outside an open chromatin region in post mortem PFC and iNeurons (**Figure 5E**), whereas the identified emVar (rs35346340) was located in the middle of a strong ATAC-Seq peak in post mortem PFC and iNeurons (**Figure 5E**). This observation suggests that distinct genetic variants within the same LD-structure contribute to gene expression changes in a highly context specific fashion.

The second emVar (rs35346340) detected at the *FURIN/FES* locus showed no allelic difference under stimulation, but exhibited lower expression for the C allele under TTX treatment (**Figure 5G** bottom) consistent only with the *FES* eQTL effect (**Figure 5F** bottom right).

The former findings indicate that stimulus dependent activity of genetic variants might be a critical mechanism contributing to the genetic basis of mental illness. In order to further investigate this hypothesis, we mapped the activity dependent gene expression programs and open chromatin landscape in iPSC derived neurons from 10 distinct donors (5 HC/5 SCZ), comparing their RNA-Seq and ATAC-Seq profiles (**Table S4)** under baseline and stimulation (KCl dependent depolarization) conditions. Consistent with previous reports, we observed widespread activity dependent open chromatin (**Figure 5H**) and gene expression remodeling (**Figure S5E**). This remodeling was heavily biased for increased accessibility/expression upon stimulation and included expression of many of the canonical activity responsive genes (**FigureS S5E,F**). We leveraged the set of stimulus responsive ATAC-Seq peaks detected across donors to test the hypothesis whether or not the landscape of activity dependent GREs is enriched for disease associated polygenic risk using partition heritability analysis ^7^. This analysis identified the set of open chromatin regions exhibiting increased accessibility upon stimulation as significantly enriched for SCZ associated polygenic risk (**Figure 5I**), further supporting the MPRA results.

In summary, these observations underscore the importance of interpreting the functional consequences of disease associated genetic variation not only under steady state conditions but also highlights the activity dependent action of SCZ associated genetic variants as a potential mechanism contributing to subtle alterations in neuronal plasticity properties.

### Identification of functional targets of emVars

In order to understand the molecular consequences of functional disease associated genetic variants, it is essential to identify their context specific target genes (Q3). Against this background, eQTL mapping and chromatin conformation capture by HiC has proven particularly useful in associating non-coding regions with putative target genes^12,27^. Therefore, we leveraged publicly available capture-HiC datasets from human post mortem PFC as well GRE-gene associations based on the ABC-contact model to identify putative target genes of the emVar set.

Using this approach, we were able to link 393 emVars to at least one putative target gene (**Figure 6A**). However, given the relatively limited resolution of currently available HiC data sets, this strategy frequently associated multiple genes or no genes to the same variants, retaining substantial ambiguity. Further integration of eQTL data from PFC ^11^ increased the fraction of variants associated to candidate genes to 85.5%, but retained frequent many-to-many associations. In particular, 74.7% of emVar genes were associated with more than one emVar (**Figure 6B**). In order to understand whether or not these statistical associations were also supported by functional connections, we implemented a GRE perturbation system in iPSC derived iNeurons similar to previous approaches^28^ (**Figure 6C**). To that end, we created a genetically modified iPSC line harboring a doxycycline-inducible catalytically inactive Cas9 fused to the Krüppel-associated box (dCas9-KRAB) in the *AAVS1* locus (**Figure S6**). This approach has been previously shown in other systems to robustly shut-down regulatory elements and map functional GRE-gene connections with relatively high throughput and amenable to parallelization^29,30^. Using this knock-in system, we first tested the hypothesis, whether or not pEEs harboring SCZ associated genetic variants had the capacity to modulate gene expression in iPSC derived neurons. For this pilot experiment, we selected the well-known SCZ risk locus TCF4 and independently targeted 6 GREs across the locus containing SCZ associated variants with or without emVar status with at least 2 gRNAs each (**Figure 6D, Table S7**). Subsequently, we evaluated the expression of TCF4 in iPSC derived neurons at day 49 by qPCR relative to a within locus control (**Figure 6D**). This experiment identified three distinct regions with the capacity to contribute to TCF4 expression in iPSC derived neurons, affecting expression between 10 and 20%. The strongest effect was observed for region B (**Figure 6D**), harboring an emVar and overlapping with open chromatin regions in human PFC and iPSC derived iNeurons. Other hits contained the TCF4 promoter of the long form (**Figure 6D**) as well as one region harboring a SCZ associated SNP classified as non-enhancer emVar (**Figure 6D** region A).

**Figure 6.**
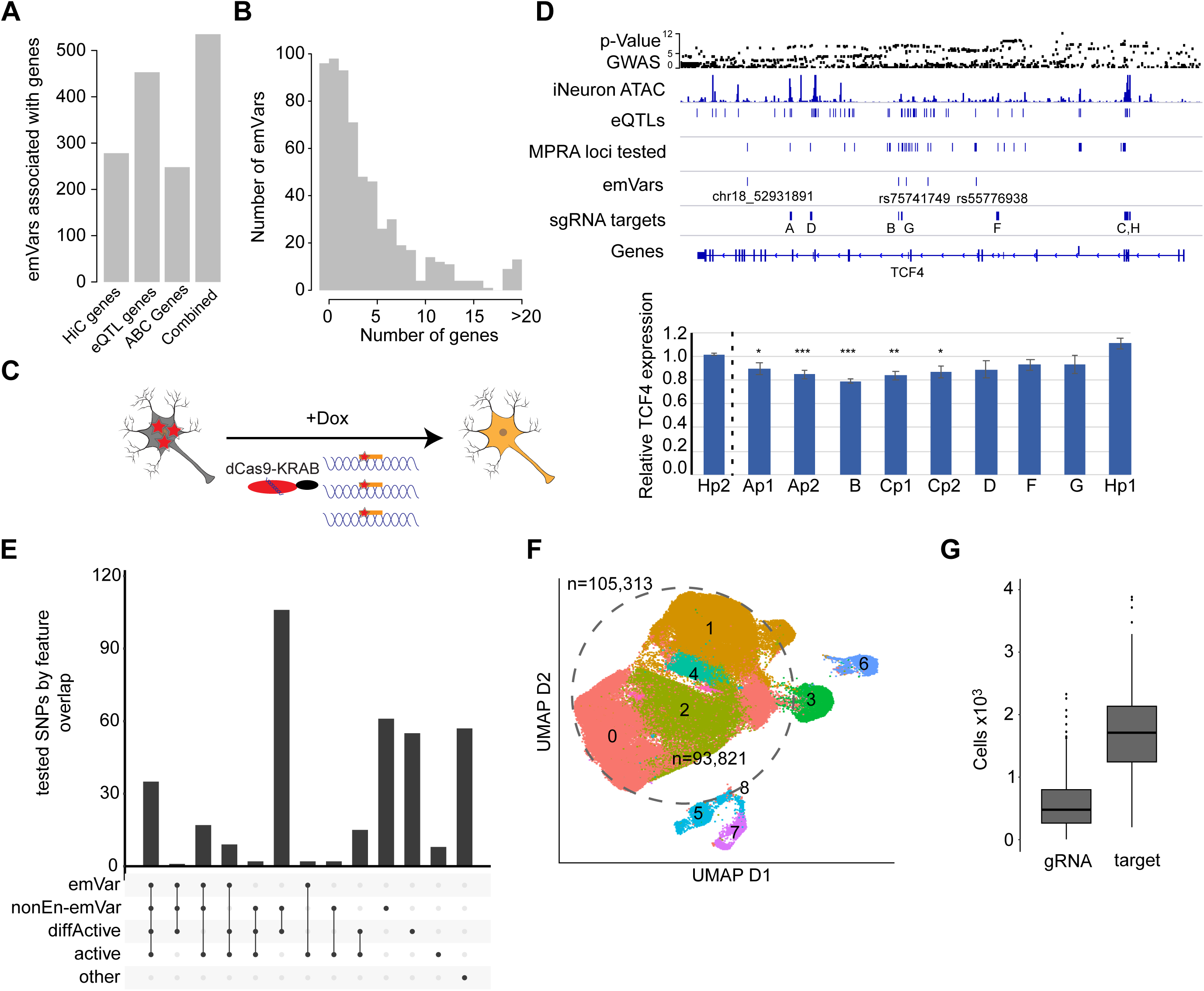
Functional interrogation of emVars. (A) Number of emVars (y-axis) associated with at least one gene by HiC in human adult post mortem PFC, eQTLs in human adult post mortem PFC, the abc model and combined. (B) Number of genes (x-axis) per emVar (y-axis) based on HiC, eQTL and the abc model combined. (C) Schematic of dCas9-KRAB experiments to silence GREs harboring SCZ associated genetic variants in iNeurons. (D) Top: IGV overview of the TCF4 locus including annotation for p-value (Ripke, 2014) - GWAS based −log10(p-value) of SCZ association, iNeuron ATAC - ATAC-Seq data iNeurons, eQTL - in human adult post mortem PFC, MPRA loci tested – all SCZ associated variants interrogated by MPRA, emVars – emVars identified across all conditions, gRNA targets – location of gRNA targets (3 gRNAs per target), letters indicate target region. Bottom: Results of dCas9-KRAB perturbation experiment in iPSC derived iNeurons at day 49. Y-axis shows fold change of the relative expression (vs MAP2) of the TCF4 gene compared in each individual perturbation experiment (x-axis) measured by qPCR. Expression was normalized to the TCF4 expression of the locus internal gRNA control H2. H1 represents an additional targeting control. Stars indicate significance of downregulation (one sided t-test) compared to H2 control with * p-value<0.05, **pvalue<0.005, ***p-value<0.0001 across 4 technical and 2 biological replicates. Error bars indicate standard error. (E) Variant classification of MPRA tested SCZ associated genetic variants (n=467) selected for interrogation by CROP-Seq: emVar – classification as emVar, non-enhancer emVar – sequence element with allele specific activity not meeting the minimal expression threshold, diffActive – differentially active element between cell types or conditions, active – element classified as active in at least one condition. F. UMAP representation of single cell CROP-Seq experiment in day 35 iPSC derived iNeurons with each dot representing a single cell. Numbers indicate cluster membership and dashed ellipsoid indicates neuronal cells. G. Distribution of number of cells per single gRNA (y-axis left) or per target (3 gRNAs per target, y-axis right) for CROP-Seq experiment. (F) UMAP representation of single cell CROP-Seq experiment in day 35 iPSC derived iNeurons with each dot representing a single cell. Numbers indicate cluster membership and dashed ellipsoid indicates neuronal cells used for further CROP-Seq analysis. (G) Distribution of number (y-axis) of cells per guide (left) or cells per target (right) for CROP-Seq experiment.

In summary, this experiment provided additional evidence for the observation of multiple EEs contributing to gene expression of the same gene. Moreover, each element can be affected by disease associated genetic variation, potentially independently modulating gene expression.

Based on these results for a single locus, we sought to identify functional targets of SCZ associated genetic variants in a more systematic fashion. To that end, we implemented a CROP-Seq based screening approach for EEs similar to previous experiments conducted in K562 cells ^30^, combining pooled CRISPR screening with single-cell RNA-Seq. Based on complexity constraints, we designed a pooled CRISPR library against 150 distinct SCZ associated genomic regions interrogated in our MPRA experiments with 3 gRNAs each (**Table S5)**. In total, we interrogated 70 emVars and 177 non-enhancer emVars from the set tested by MPRA. Most of these variants were annotated as eQTLs and overlapped with epigenomic signatures of enhancer elements across multiple primary and in vitro derived cell and tissue types (**Figure 6E**, **Figure S7A**).

Subsequent to gRNA pool complexity validation (**Figure S7B**), we triple infected iPSC/iNeurons with the gRNA pool at iPSC and neuronal stages to maximize gRNA copy number per cell and performed scRNA-Seq of six distinct iNeuron pools at day 35 of differentiation. This experiment resulted in 105,313 usable cells (**Methods**), distributed across 8 distinct cell clusters (**Figure 6F**), representing mostly excitatory neurons (**Figure S7C**). Across these cells, we obtained a median coverage of 479 cells per guide and a median of 1,711 cells per non-coding target region (collapsing all 3 guides per target **Figure 6G**). To not confounding our analysis by strong cellular heterogeneity, we restricted all subsequent analyses to the neuronal clusters 0-2 and 4 (**Figure 6F**, dashed ellipsoid, **Figure S6C**), leaving 93,821 cells.

In order to pinpoint the target genes of emVars, we next identified gRNA pools (collapsing the 3 gRNAs per target) that were associated with a significant reduction in potential target gene expression. This analysis revealed an enrichment of significant associations in the enhancer targeting gRNAs pools compared complexity matched permuted control sets and non-targeting controls (**Figure 7A**). Overall, this strategy identified 40 (FDR ≤0.1) unique gRNA-gene pairs distributed across 34 distinct gRNA pools (25 targeting MPRA non-coding regions) that we defined as hits as well as 85 gRNA-gene pairs at nominal significance (p-value≤0.05, 66 unique gRNA pools, 56 targeting MPRA non-coding regions) defined as nominal hits (**Figure 7B**). The gRNA hit pools reduced the expression of 32 (61 for tentative hits) unique genes by 10% on average (**Figure 7C**), with the majority of gRNAs pools affecting a single gene (**Figure 7D, Table S5**). Most significant gRNA-coding gene pools harbored either an emVar (36%/35.7) or a non-enhancer emVar (36%/41) in close vicinity and overlapped with an eQTL in adult PFC as well as an ATAC-Seq peak in iNeurons (**Figure S7D**).

**Figure 7.**
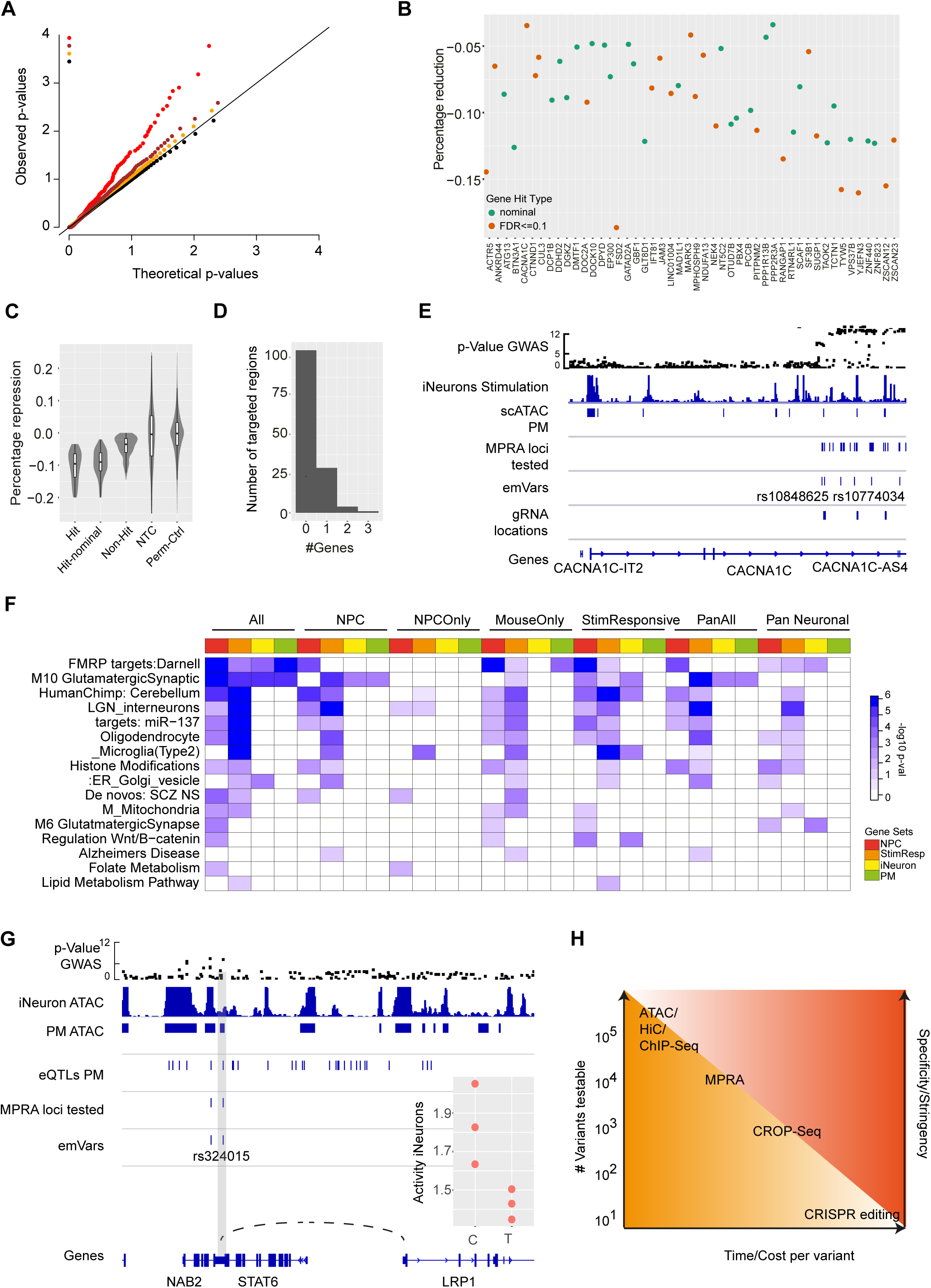
Functional characterization of emVars by CROP-Seq. (A) Quantile-quantile plot of differential expression tests showing the distribution of observed vs. expected p-values targeting gRNA pools (red, n=181), permuted control gRNA pools of matching size and complexity distribution (orange, n=500), non-targeting controls (n=9) and bootstrapped p-value for randomly sampled cell sets of equal size and complexity (black). P-values originate from one sided t-tests. (B) Summary of CROP-Seq screen results showing all down-regulated target genes (x-axis) along with the percentage of downregulation (y-axis) by individual gRNA pools (dots) for significant gRNA pool – gene hit pairs based on FDR (FDR ≤0.1, orange) or nominal significance (p-value ≤0.05, green). (C) Effect size distribution in terms of percentage down-regulation (y-axis) of significant gRNA-gene pairs based on an FDR ≤0.1, nominal significance (p-value ≤0.05), non-significant gRNA-gene pairs (Non-Hit) and filtered NTC-gene pairs (NTC) and the permutation control gRNA-gene pair set (Perm Ctrl, x-axis). (D) For each gRNA pool (y-axis) number of genes downregulated (x-axis). (E) Example IGV overview of two significant gRNA-gene pairs at the CACNA1C locus. Grey boxes indicate gRNA locations associated with significant downregulation of the CACNA1C gene. (F) Selected results of pathway enrichment analysis (x-axis) for genes linked to distinct emVar Sets (y-axis) by HiC, eQTL or CROP-Seq and expressed in the tissue indicated by the color code shown on the left. Pathway enrichment results are shown as −log_10_ p-value from Fisher’s exact test only for pathways with an FDR≤0.1 and capped at 6. For all pathway enrichment results see Supplementary Table 6. (G) Left: IGV overview of the LRP1 locus including annotation for p-value (Ripke, 2014) - GWAS based −log10(p-value) of SCZ association, iNeuron ATAC - ATAC-Seq data iNeurons, PM ATAC – ATAC-Seq data from human post mortem brain, eQTL - in human adult post mortem PFC, MPRA loci tested – all SCZ associated variants interrogated by MPRA, emVars – emVars identified across all conditions. Right: Allele specific GRE activity for the C and T allele of the rs324015) SNP based MPRA measurements in iNeurons. Each dot represents an independent MPRA assay replicate. (H) Schematic summarizing the characteristics of different functional genomic assays with respect to the number of genetic variants that can be interrogated (y-axis left), the time/cost required for each assay (x-axis) and the specificity/stringency of the obtained results (y-axis left).

This included emVars at many well-known SCZ risk genes (**Figure 7B**) such as the *CACNA1C* gene (**Figure 7E**). At this locus, we identified two distinct emVars within the intronic region of the gene that overlapped or were in close vicinity (7kb) of two distinct gRNA pool landing sites, 33kb apart from each other. Importantly, the emVar rs12315711 was directly adjacent to a previously identified functional genetic variant with strong evidence for *CACNA1C* promoter interaction (rs2159100) based on experiments in HEK cells^31^. However, while the rs2159100 variant did also show highly significant allele specific activity in primary mouse cultures (FDR 4.89 10^−08^), it was not classified as an emVar due to the overall low baseline activity of the surrounding candidate regulatory element (non-enhancer emVar), consistent with the absence of ATAC-Seq signal in iNeurons. In addition, the CROP-Seq screen identified emVars, such as a functional SNP in the intronic region of the *KCTD13* gene that was connected to the promoter of the *TAOK2*, 54kb away with (**Figure S7E**).

In summary, these experiments enable the functional association of emVars to their target genes, highlighting the power of this approach to narrow down the number of putatively causal disease associated genetic variants impacting gene expression.

Moreover, these experiments suggest that only a very small fraction of candidate eQTLs and disease associated variants have the potential to contribute to changes in gene expression. Based on these results, it is important to note that the functional non-coding genetic variants do not necessarily overlap with the most highly disease associated genetic variant and can exhibit an association strength below genome-wide significance.

### Biological insights into context specific cellular processes modulated by functional SCZ associated NCV

Finally, we leveraged the combined results of the MVAP pipeline to refine the current interpretation of genetic SCZ risk factors (Q4), including many sub-threshold genetic variants. To that end, we performed cell type specific pathway enrichment analyses of all genes associated with distinct emVars based on the CROP-Seq screen, HiC or eQTL data. This analysis confirmed the major biological themes previously found by genetic analyses, including immune, histone modification and synapse biology related processes (**Table S7, Figure 7F**). This analysis also identified the glutamatergic synapse enriched in emVar associated genes in PFC and iNeurons as well as genes enriched in CNVs overrepresented in SCZ patients (**Figure 7F**). Interestingly, the set of pan-neural emVars was also associated with genes involved in synaptic signaling in autism spectrum disorder. Moreover, we detected an enrichment of the AGE/RAGE pathway and genes involved in lipid and cholesterol metabolism in the set of stimulus responsive emVar genes (**Figure 7F**) providing additional evidence for a potential role of these processes in SCZ pathophysiology affecting neuronal cells. More specifically, we identified multiple functional SCZ associated NCVs modulating key genes in involved in lipid homeostasis and signaling such as the Lysophosphatidic Acid Receptor 2 (LPAR2) and the LDL Receptor Related Protein 1 (LRP1). For the former, we identified functional NCVs located within an iNeuron ATAC-Seq peak located in an intron of the YJEFN3 gene ~100kb upstream of the LPAR2 promoter. Similarly, we identified a NCDV modulating *LRP1* expression, located within an ATAC-Peak in the 3’UTR of the *STAT6* gene 35kb upstream of the *LRP1* promoter (**Fig. 7G**). Interestingly, both variants have recently passed the genome wide significance threshold for SCZ association based on the latest GWAS. In this context, LRP1 is of particular interest, as it is cleaved by FURIN^32^, another SCZ associated gene modulated by NCDVs. LRP1 is involved in a diverse array of signaling and other cellular processes such as inflammation ^33^ and is expressed in various tissues, including the brain. In the latter context, LRP1 has been implicated in both development, maturation and homeostasis of neuronal cells ^34^. Moreover, it has recently been implicated a key gene in Alzheimer disease, serving as a master regulator of tau uptake and spread by controlling the endocytosis of TAU/MAPT^35^.

In summary, these observations illustrate the added value of the results provided by MVAP analysis to identify candidate biological processes associated with SCZ down to the functional genes and genetic variants that drive these associations. Thus, these insights provide a valuable starting point for in depth future studies on the molecular, cellular and circuit level role of SCZ associated NCV.

## Discussion

The results presented in the previous sections constitute a significant step towards addressing four key questions in the field of SCZ genetics and provide a substantially refined list of functional SCZ associated SNPs, that is likely highly enriched for causal genetic variants. The strategy outlined here leveraged well defined molecular traits (gene-expression) to narrow down the number of genetic variants functional on a molecular level. This approach is enabled by sequentially arraying experimental techniques with decreasing throughput/comprehensiveness, but increasing stringency and specificity, starting from (1) ChIP-Seq/ATAC-Seq/HiC, over (2) MPRA, and (3) CROP-Seq towards individual variant interrogations in their endogenous context by CRISPRi-qPCR (**Figure 7H**). The coherent integration of these assays into a massively parallel variant annotation pipeline empowered the identification of hundreds of highly cell type and condition specific SCZ associated functional emVars, out of thousands of statistically associated variants. Our results on cell and condition specificity of these results clearly underscore the need to analyze SCZ associated NCV function in disease relevant cell types of the CNS.

This map of functional variants and their context specific action in disease relevant cell types constitutes a unique resource for future mechanistic studies. In particular, this resource provides a starting point to select the appropriate variant and cellular context to characterize its cellular effects in detail, directly enabling focused mechanistic studies to obtain insights into associated molecular pathomechanisms.

Our results also indicate that the statistical association strength of a GWAS locus is not related to the likelihood of the presence of an emVar. However, the latter is also not expected considering that it is unlikely that disease associated regions of the genome harbor more regulatory elements susceptible to genetic variation. Instead, the latter is more likely to be a function of gene density and the complexity of the cis-regulatory architecture of each locus. This complexity is illustrated by the MPRA based findings at the *FURIN/FES* locus, suggesting that a single SNP might have different effects on different genes depending on the stimulation condition.

These observations render the cell type specific functional association of emVars with target genes even more important, highlighting the value of the extension of current cell type/condition specific annotations of SCZ emVars. In this context, our results in NPCs and under various stimulation paradigms represent the first assessment of SCZ associated genetic variant action under these conditions, revealing the enrichment of stimulus responsive open chromatin regions for SCZ polygenic signal. These results also highlight the limitations of current epigenomic and eQTL based annotation and selection strategies for SCZ NCVs, as only 12.3% of NCVs overlapping with these annotations, were deemed functional, underscoring the added value of the MVAP pipeline.

Overall, our results support previous biological interpretations of SCZ GWAS results. However, now tying them to specific SNP-gene combinations in individual cell types and conditions, a substantial step beyond the current state of the art. Moreover, our results suggest multiple additional relevant biological processes that are likely subject to modulation by emVars in SCZ such as lipid metabolism and signaling in neurons, providing also the basis of these associations in terms of functional gene-variant connections.

However, the approach presented here is inherently limited to genetic variants that exert their effects through the modulation of gene expression levels, which are currently estimated to apply to up to 50% ^36^ of disease associated SNPs. In addition, the MPRA variant employed here is not capable of interrogating GREs with silencing function, actively contributing to the reduction of gene expression levels^37^. Thus, the current results likely underestimate the number of genetic variants capable of modulating gene epxression levels. Moreover, genetic variants affecting other processes such as splicing, mRNA stability, mRNA/protein localization or protein function are not captured. For several of the latter, other high throughput approaches have been/are developed that can eventually provide a comprehensive functional genomic toolbox to dissect NCDVs at scale in disease relevant cell types^38–40^.

Going beyond SCZ, the integrated MVAP approach presented here provides a general strategy to functionally annotate, fine map, and characterize NCDVs in a massively parallel fashion. This rationale provides one approach to address the large and vastly increasing number of statistically disease associated NCVs with individual effect sizes barely measurable with respect to physiological or even cellular traits.

In summary, the application of the MVAP pipeline yields a disease associated genetic variant set highly enriched for functional variants that can in the next step be leveraged for functional follow up experiments through multiplexed genome editing or combinatorial modeling studies of polygenic, eQTL mediated effects.

## Acknowledgments

We would like to thank the members of the Department of Translational Psychiatry at the MPIP and in particular Elisabeth Binder, Monika Rex-Haffner and Dietmar Spengler for their comprehensive support and critical discussion throughout the project.

## Funding

This work was supported by grants from the BMBF eMed program grant 01ZX1504 to M.J.Z. and the Max-Planck-Society.

This study used data from the CommonMind consortium provided through NIMH. Data for this publication were obtained from NIMH Repository & Genomics Resource, a centralized national biorepository for genetic studies of psychiatric disorders. Data were generated as part of the CommonMind Consortium supported by funding from Takeda Pharmaceuticals Company Limited, F. Hoffman-La Roche Ltd and NIH grants R01MH085542, R01MH093725, P50MH066392, P50MH080405, R01MH097276, RO1-MH-075916, P50M096891, P50MH084053S1, R37MH057881, AG02219, AG05138, MH06692, R01MH110921, R01MH109677, R01MH109897, U01MH103392, and contract HHSN271201300031C through IRP NIMH. Brain tissue for the study was obtained from the following brain bank collections: the Mount Sinai NIH Brain and Tissue Repository, the University of Pennsylvania Alzheimer’s Disease Core Center, the University of Pittsburgh NeuroBioBank and Brain and Tissue Repositories, and the NIMH Human Brain Collection Core. CMC Leadership: Panos Roussos, Joseph Buxbaum, Andrew Chess, Schahram Akbarian, Vahram Haroutunian (Icahn School of Medicine at Mount Sinai), Bernie Devlin, David Lewis (University of Pittsburgh), Raquel Gur, Chang-Gyu Hahn (University of Pennsylvania), Enrico Domenici (University of Trento), Mette A. Peters, Solveig Sieberts (Sage Bionetworks), Thomas Lehner, Stefano Marenco, Barbara K. Lipska (NIMH).

## Author Contributions

CR performed MPRA, ATAC-Seq, RNA-Seq, CRISPRi for TCF4 and iPSC related experiments. MG generated dCas9 knock-in iPSC lines, performed the CROP-Seq experiment and performed qPCR data analysis. AHe performed the MPRA experiments in primary mouse cultures guided by MR. RA performed all NPC experiments, AH and SM assisted with iPSC cell culture and performed RNA-Seq experiments. CF assisted with iPSC cultures and TCF4 characterization experiments. VM, LW and LT assisted with primary data processing. MJZ conceived the study, performed data analysis, supervised the study and wrote the manuscript with contributions from all authors.

## Supplementary Figures

**Figure S1.**
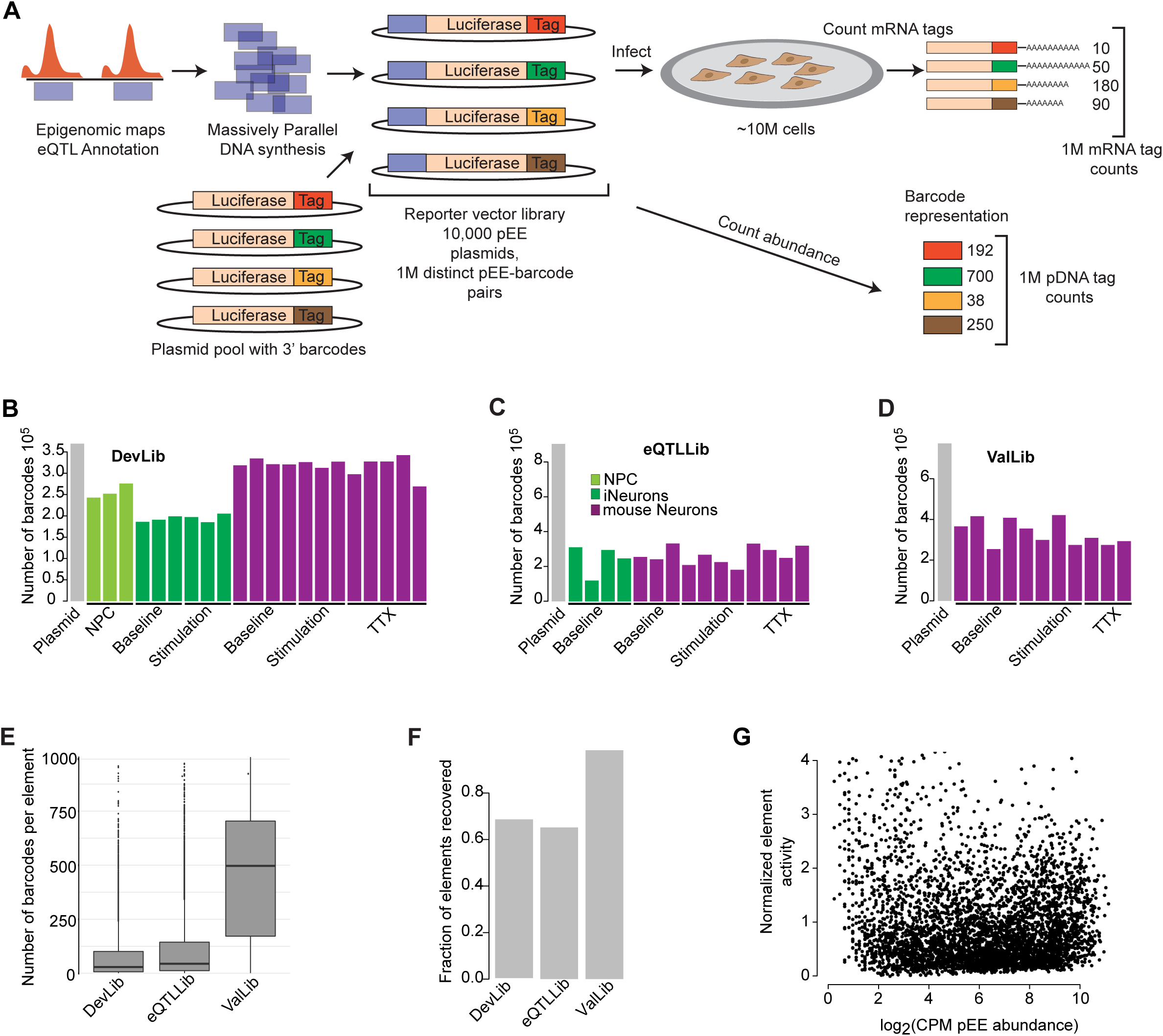
MPRA quality metrics. (A) Schematic representation of MPRA workflow. (B-D) Total number of unique barcodes detected in the plasmid library (grey bar), as well as in mRNA obtained from the individual experimental MPRA conditions for the developmental (DevLib), eQTL (eQTLLib) and validation library (ValLib) in the different neural cell types: NPCs (light green), iNeurons (dark green), and mouse primary neurons (purple). (E) Distribution of the number of individual barcodes per tested sequence element (each allele is counted separately). Boxes indicate 25^th^ and 75^th^ percentile and whiskers indicate 1.5 interquartile range from the hinge. (F) Fraction of individual sequences contained in each library design that were recovered at quantifiable levels from the RNA sequencing of each library. (G) Distribution of sequence element activity (x-axis) normalized by sequence abundance in plasmid library (x-axis) as a function of total element abundance in the original plasmid library for the DevLib in iNeurons.

**Figure S2.**
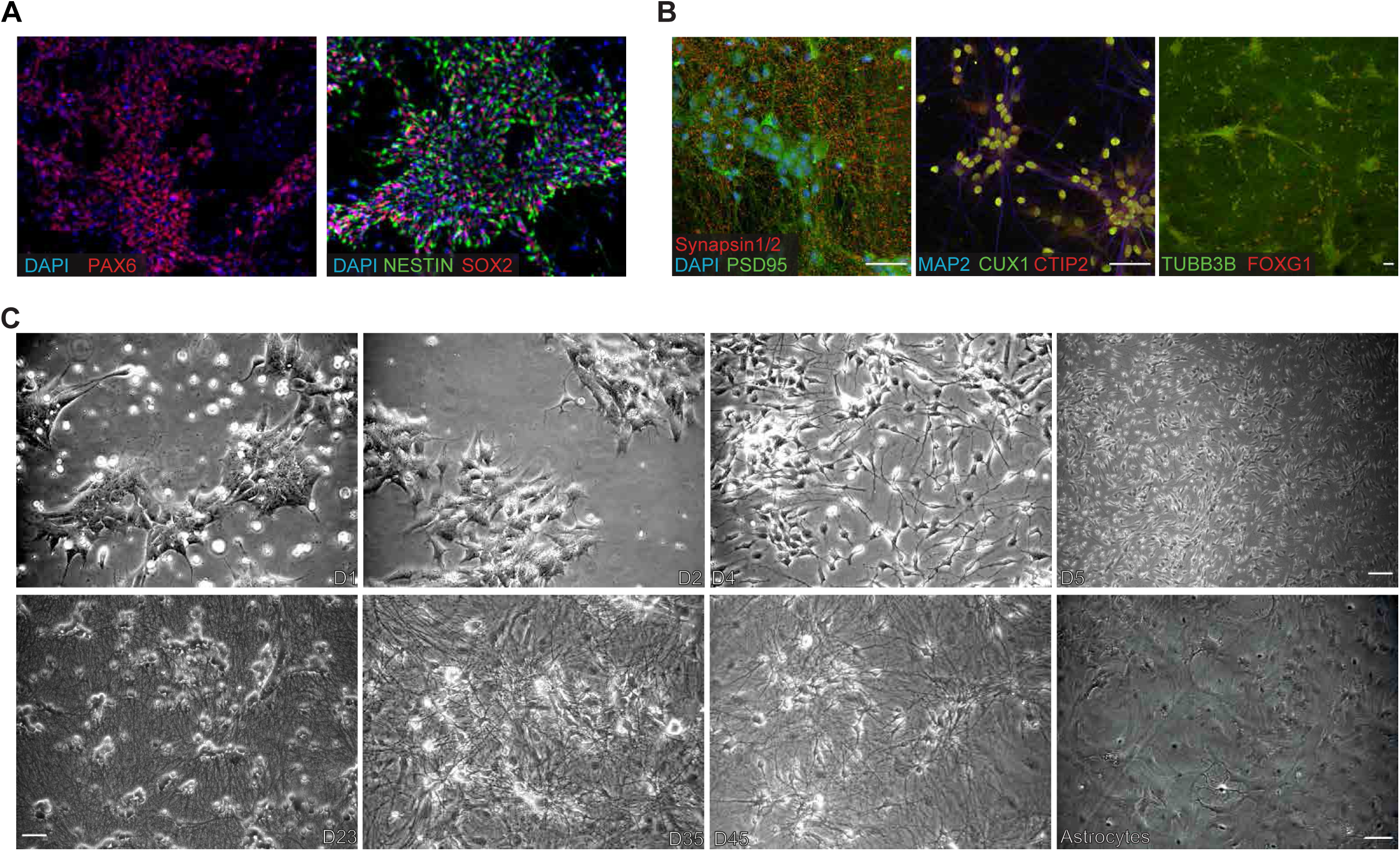
Characterization of in vitro model system. (A) Immunohistochemistry of NPCs at day6 of differentiation from iPSCs. DAPI is used as counterstain for DNA located in the nucleus of the cells. NPCs are positive for PAX6 (forebrain NPC marker), NESTIN (immature NPC neurite marker), and SOX2 (immature neural marker). (B) iNeurons on day49 are expressing cortical layer marker CUX1 and CTIP2. Neurons have a dense neuronal network indicated by the neurite marker TUBB3B (TUJ1, β3-Tubulin). Alongside the neurite network, the excitatory presynaptic marker SYNAPSIN1/2 and postsynaptic marker PSD95 can be located. Scale bar indicates 50 µm. (C) Phase contrast images of different stages during the iNeuron differentiation. Image on the bottom right depicts the primary mouse astrocytes used for the maturation of the iNeurons. Scale bar indicates 50 µm.

**Figure S3.**
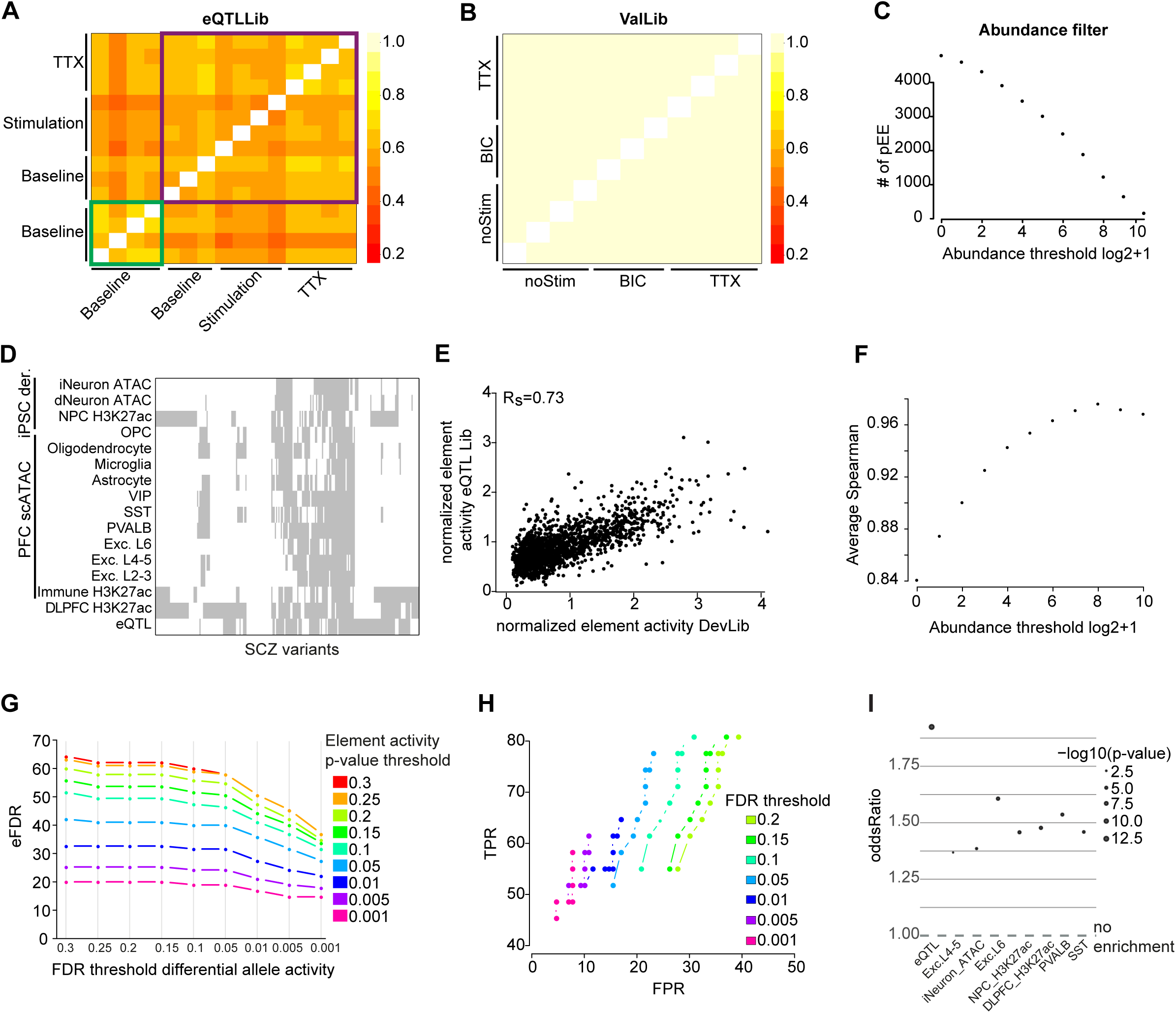
MPRA quality metrics and threshold calibration. (A-B) Spearman correlation heatmap of cpm normalized log2 ratio between the MPRA mRNA counts and their abundance in the original plasmid library for all pEE measured in all conditions of the eQTL or Validation library. Green box signifies correlation of MRPA results from human iNeurons, purple box demarcates results for MPRA experiments in mouse primary neurons. (C) Number of recovered sequence elements in the eQTL lib (y-axis) as a function of cpm normalized element abundance in the original plasmid library (x-axis). (D) Annotation of all SCZ associated genetic variants selected for MPRA testing in the ValLib (x-axis) with various epigenomic annotations (y-axis) derived from iPSC derived neurons (iNeuron, dNeuron, neural precursor cells (NPCs)), single cell ATAC-Seq of adult human post-mortem tissue from the prefrontal cortex (PFC), H3K27ac ChIP-Seq profiles across immune cell types and bulk PFC tissue as well as eQTLs detected in PFC. (E) Consistency of allele specific pEE activity between the eQTLib and the DevLib (n=193) in mouse primary neuronal cultures at baseline conditions. (F) Average Spearman correlation across replicates of DevLib (y-axis) as a function of minimal element abundance in the plasmid library (x-axis). (G) Empirical false discovery rate (eFDR, y-axis) as a function of the difference between the two alleles of each sequence element and the minimal element activity threshold (x-axis) defined as the most significant activity across both alleles (colored lines) for 95 random control SNPs in the validation library. (H) True positive (y-axis) and false positive rate (x-axis) for allele specific SNP activity measured in the DevLib in primary mouse cultures (baseline condition) using the results of the validation library as ground truth. Results are shown as a function of differential allele FDR thresholds (dots, from highest dot to lowest 0.3, 0.25,0.2, 0.15, 0.1, 0.05, 0.01, 0.005, 0.001) and minimal element activity threshold (colored lines). (I) Enrichment of sequence elements identified as active (p-value≤0.005 compared to the negative control elements) compared to non-active elements based on their overlap with eQTL, ATAC-Seq or H3K27ac peaks in various cell types and tissues (x-axis, eQTL – eQTL (FDR≤0.05) in adult human post mortem PFC, single cell ATAC-Seq in PFC: Excl.4-5 – excitatory neurons layer 4-5, Exc. L6 – excitatory neurons layer 6, PVALB-parvalbuminergic interneurons, SS T-somatostatin interneurons, PFC H3K27ac – ChIP-Seq in dorso lateral prefrontal cortex of adult human post mortem brain, iNeuron – ATAC-Seq iPSC derived neurons at day 49 of culture). y-axis indicates odds ratio and point size indicates p-value of Fisher’s-exact test results.

**Figure S4.**
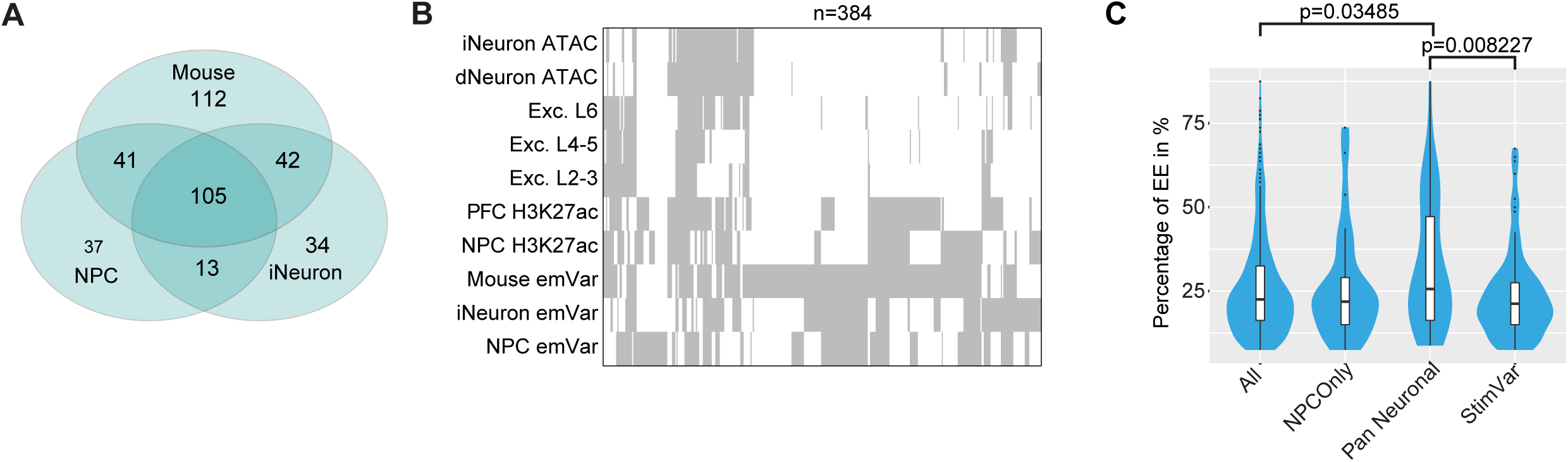
DevLib MPRA results. (A) Overlap of emVars in the DevLib detected in primary mouse cortical neurons under baseline conditions, iPSC derived NPCs and iPSC derived iNeurons. (B) Genomic feature annotation (x-axis) of emVars detected in mouse, iNeurons or NPCs (y-axis). Annotation include NPC H3K27ac – ChIP-eq for H3K27ac in iPSC derived NPCs, DLPFC H3K27ac – ChIP-Seq in dorso lateral prefrontal cortex of adult human post mortem brain, single cell ATAC-Seq in PFC: Exc. L2-3 – excitatory neurons layer 2-3, Exc. L4-5 – excitatory neurons layer 4-5, Exc. L6 – excitatory neurons layer 6, PVALB-parvalbuminergic interneurons, SST – somatostatin interneurons, iNeuron/dNeuron – ATAC-Seq iPSC derived iNeurons at day 49 of culture or Qi Neurons (dNeurons) at day 63 of culture. (C) Percentage of EE regions harboring emVars (x-axis) with H3K27ac signal across 80 diverse human cell and tissue types (y-axis).

**Figure S5.**
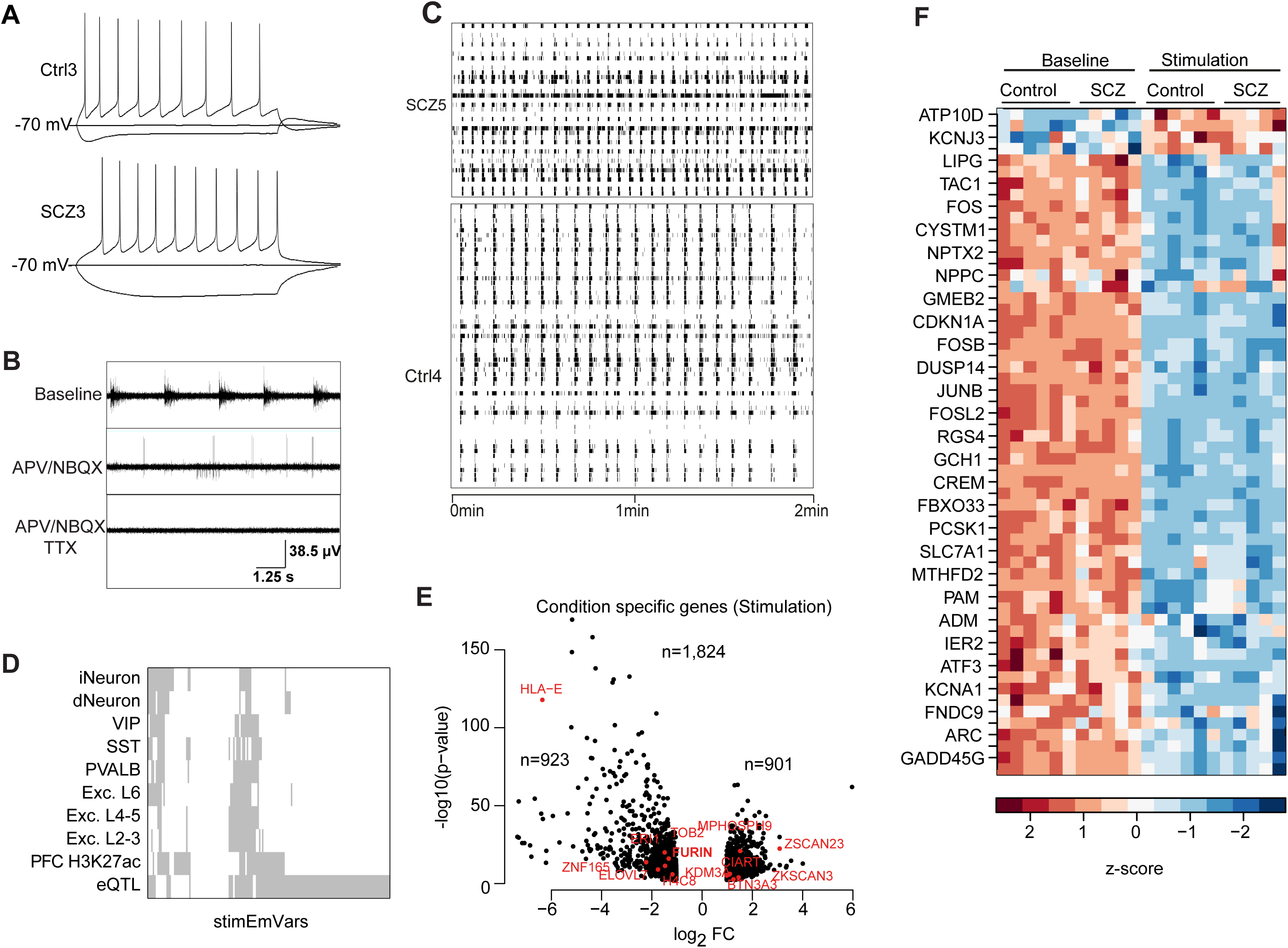
Electrophysiological characterization of iPSC derived neurons. (A) Representative single cell whole cell current clamp analysis results of iPSC derived iNeurons at day 45-51 of differentiation from healthy donor (top, C3) or SCZ patient (bottom, S3). (B) Representative multi-electrode-array field recordings of iPSC derived neurons under baseline conditions (top), after addition of APV and NBQX (middle) and APV, NBQX, and TTX (bottom) over 10 seconds. (C) Representative multi-electrode-array recordings of iPSC derived iNeurons at day 49 from a healthy donor (C6) and a SCZ donor (S4) over 2 min (x-axis). Y-axis shows all active electrodes with black lines indicating detected action potentials. (D) Genomic feature annotation (y-axis) of stimEmVars ––(x-axis). Annotation include eQTL – eQTL in adult human post mortem PFC, DLPFC H3K27ac – ChIP-Seq in dorso lateral prefrontal cortex of adult human post mortem brain, single cell ATAC-Seq in PFC: Exc. L2-3 – excitatory neurons layer 2-3, Exc. L4-5 – excitatory neurons layer 4-5, Exc. L6 – excitatory neurons layer 6, PVALB – parvalbuminergic interneurons, SST-somatostatin interneurons, iNeuron/dNeuron – ATAC-Seq iPSC derived neurons (E) Differential gene expression between baseline and KCl treated (Stimulation condition) iPSC derived iNeurons at day 49 measured RNA-Seq across n=10 distinct donors per treatment. Results are shown by −log10 p-value (y-axis) and log2 fold change per gene, where only significant genes are (FDR≤0.01, log2(FC)≥1) are shown. Negative values indicate higher expression upon stimulation. Genes highlighted in red indicate SCZ risk genes based on Ripke et al. 2014. (F) Heatmap of z-score normalized gene-expression for well-known activity regulated genes based on Tussowski et al. (2018) differentially expressed upon stimulation (KCl treatment) across iPSC derived neurons from healthy donors and SCZ individuals.

**Figure S6.**
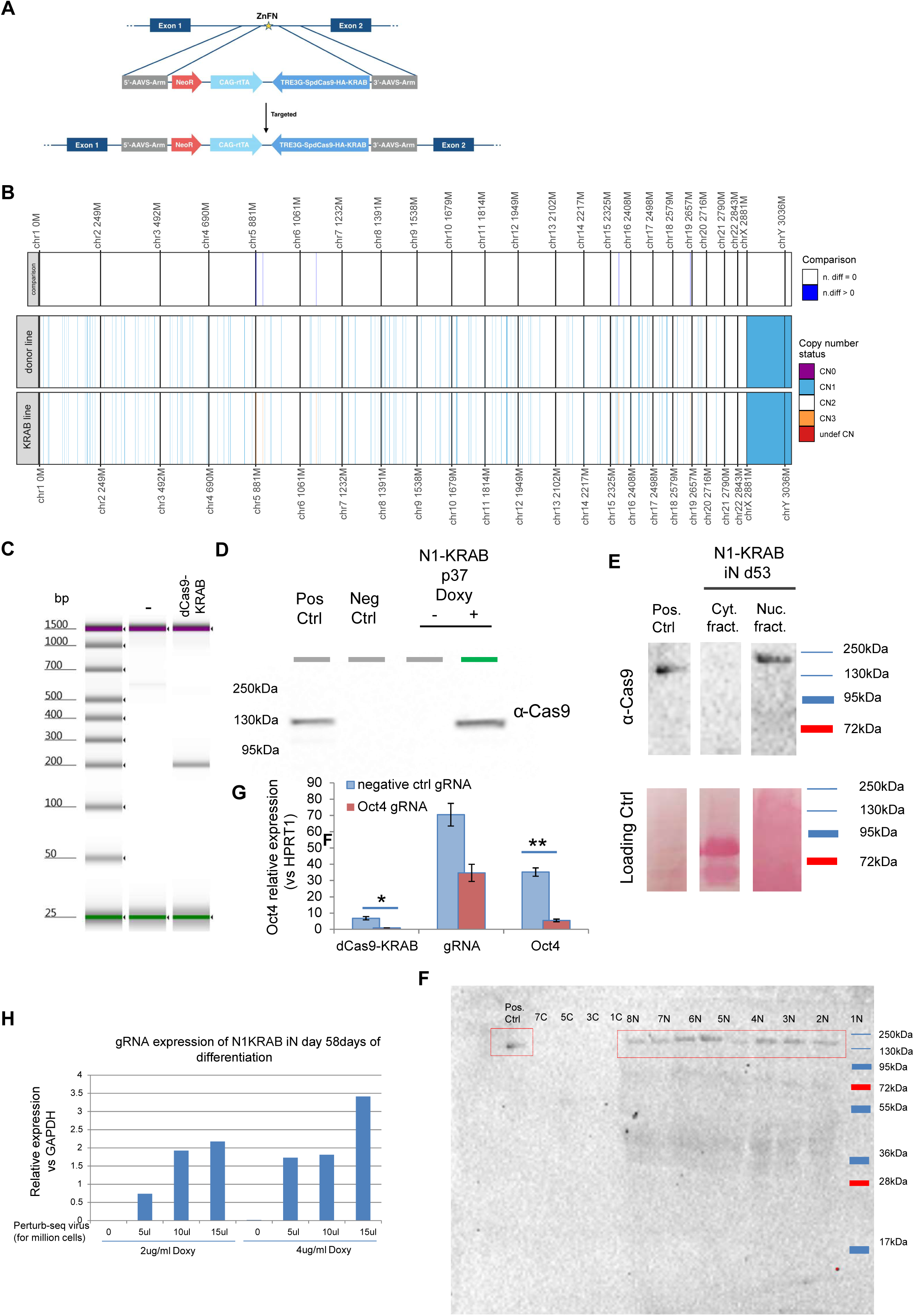
Characterization of dCas9-KRAB iPSC line. (A) Targeting strategy for dCas9-effector knock-in at the AAVS1 locus in human iPSCs (B) Karyotyping of Ctrl5 iPSC line comparing the dCas9-KRAB knock-in line to the original donor. (C) PCR validation of knock-in fragment in AAVS1 locus (D) PCR validation of dCas9 transcript induction upon doxycycline treatment in iPSCs. (E) Western blot validation of dCas9 protein expression in iPSC derived neurons (day 53) for cytosolic and nuclear fraction separately. Loading control shown below. (F) Full blot from (E): Nuclear extracts are indicated with “N”, cytoplasmatic extracts with “C”. The positive control is a whole cell extract from N1KRAB iPSCs treated for 14 days with DOXY 2ug/ml. N1KRAB iN day61 of differentiation : 1- Ctrl (not infected with CROP-seq virus) [2ug/ml Doxy] 2- Ctrl (not infected with CROP-seq virus) [4ug/ml Doxy] 3- 5 µl first infection than infected 2 times with 10ul/million cells [2ug/ml Doxy] 4- 5 µl first infection than infected 2 times with 10ul/million cells [4ug/ml Doxy] 5- 10 µl first infection than infected 2 times with 10ul/million cells [2ug/ml Doxy] 6- 10 µl first infection than infected 2 times with 10ul/million cells [4ug/ml Doxy] 7- 15 µl first infection than infected 2 times with 10ul/million cells [2ug/ml Doxy] 8- 15 µl first infection than infected 2 times with 10ul/million cells [4ug/ml Doxy] (G) dCas9 expression by qPCR relative to GAPDH in day 53 iPSC derived neurons for different dox-treatment dosages and times during differentiation. (H) Functional validation using knockdown of OCT4 relative to HPRT1 expression by dCas9 in iPSC lines.

**Figure S7.**
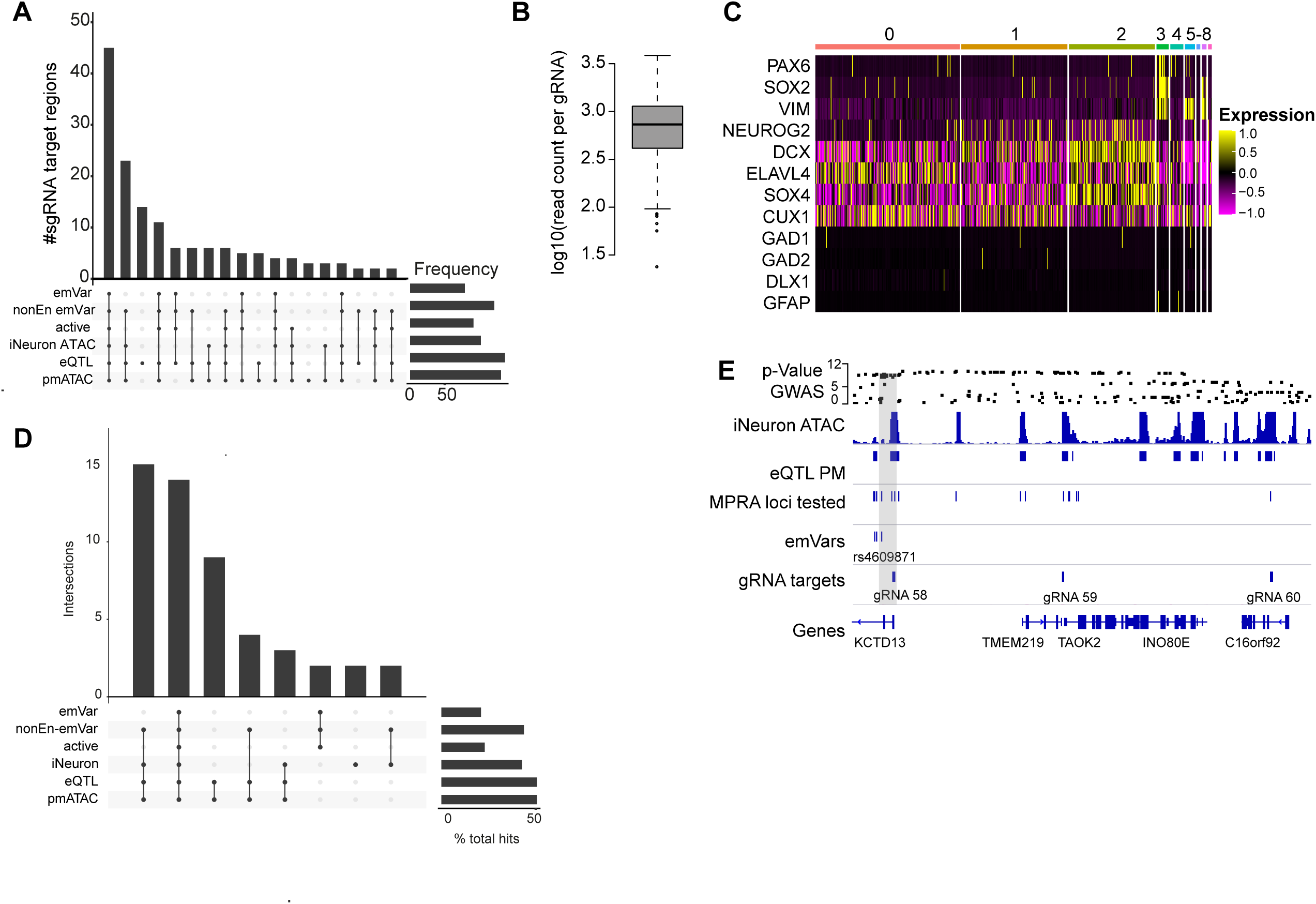
Characterization of CROP-Seq pool and experiment. (A) Annotation of gRNA pool targets (n=146) contained in the CROP-Seq library with SCZ variant classes and genomic features: emVar – classification as emVar, non-enhancer emVar – sequence element with allele specific activity not meeting the minimal expression threshold, active – element classified as active in at least one condition, iNeuron-ATAC-Seq peaks in iPSC derived iNeurons, eQTL – eQTLs measured in human adult PFC and pmATAC - open chromatin regions measured by scATAC-Seq in human post mortem cortex. (B) Normalized read count distribution of all gRNAs contained in the CROP-Seq library, recovering 100% of synthesized gRNAs. (C) Marker gene expression (y-axis) across subsampled (n=5,000) cells from the CROP-Seq experiment. Numbers on top indicate cluster ID. (D) Annotation of gRNA pool targets with a significant repressive effect on at least one target gene in the CROP-Seq screen, similar to (A). (E) Example IGV overview of one significant gRNA-gene pair at the TAOK2 locus. Grey boxes indicate gRNA locations associated with significant downregulation of the TAOK2 gene.

**Table S1: Dataset overview and basic QC stats**

**Table S2: MPRA and CROP-Seq sequences**

**Table S3: Annotated MPRA results**

**Table S4: Results of stimulation experiments in gene expression and open chromatin profiles in iNeurons**

**Table S5: CROP-Seq results**

**Table S6: Pathway enrichment results**

**Table S7: Details on resources used for molecular biology experiments**

## Notes

### Competing Interest Statement

MJR is shareholder and consultant of Systasy Bioscence GmbH. All other authors declare that they have no competing interests.

